# Serum and Tear Autoantibodies from NOD and NOR Mice as Potential Diagnostic Indicators of Local and Systemic Inflammation in Sjögren’s Disease

**DOI:** 10.1101/2024.10.24.619993

**Authors:** Shruti Singh Kakan, Sara Abdelhamid, Yaping Ju, J Andrew MacKay, Maria C. Edman, Indu Raman, Chengsong Zhu, Prithvi Raj, Sarah F Hamm-Alvarez

## Abstract

**Background:** Sjögren’s Disease (SjD) is an autoimmune disease characterized by lymphocytic infiltration of salivary and lacrimal glands (LG). The LG produces the protein-rich aqueous component of tears, and SjD-associated autoimmune dacryoadenitis (AD) may thus alter tear autoantibody composition.

**Methods:** The presence of tertiary lymphoid structures (TLS) in LG from two murine models of SjD-associated AD, male NOD and male NOR mice, were evaluated using immunofluorescence. IgG and IgA reactivity in serum and tears from these models were probed in three studies against a panel of 80-120 autoantigens using autoantibody microarrays relative to serum and tears from healthy male BALB/c mice. Data were analyzed by R package Limma.

**Results:** Analysis of immunofluorescence in LG sections from both SjD models showed TLS. Only one autoantibody was significantly elevated in tears and serum in both SjD models across all studies. Three autoantibodies were significantly elevated in serum but not in tears in both SjD models across all studies. Conversely, six IgG and thirteen IgA autoantibodies (6 sharing the same autoantigen) were significantly elevated in tears but not serum in both SjD models.

**Conclusion:** NOD and NOR mice with SjD-associated AD have distinct autoantibody profiles in tears and serum. Tear IgA isotype autoantibodies showed a greater diversity than tear IgG autoantibodies. TLS observed in LG are a likely source of the tear autoantibodies.

## Introduction

Sjögren’s Disease (SjD) is characterized by autoimmune exocrinopathy of lacrimal glands (LG) and salivary glands (SG), resulting in severe dryness of mucosal surfaces(1). This leads to clinical symptoms of dry eye and dry mouth, respectively. Extraglandular manifestations including fatigue, musculoskeletal pain, arthralgia, interstitial lung disease, and tubulointerstitial nephritis frequently occur. SjD patients also have a significantly higher risk of developing B-cell lymphoma(2).

Pathogenesis of SjD is complex and includes genetic predisposition, environmental triggers, and activation of innate and adaptive immunity(3–5). Inflammation of LG and SG via infiltrating immune cells result in the activation of apoptotic pathways in glandular epithelial cells, releasing autoantigens such as ribonucleoprotein complexes Ro/SSA and La/SSB, which further recruit dendritic cells to the exocrine glands and lead to a cycle of inflammation(2). Detection of Ro/SSA autoantibodies in serum is part of the clinical diagnosis of SjD(6). Although the presence of serum SSA autoantibody is weighted equally to a positive SG biopsy in the current American-European consensus group (AECG) classification criteria for SjD(7), SSA is not specific to SjD and is elevated in other systemic autoimmune diseases(8–10).

Tear autoantibodies may have potential as diagnostic tools denoting ocular involvement in SjD. It is established that tertiary lymphoid structures (TLS) form in the LG and SG in SjD(11). These localized TLS can generate autoantibodies distinct from those found systemically in disease. As the LG is a target organ in SjD, with autoimmune dacryoadenitis (AD) leading to changes in tear composition and volume, it is possible that the presence of specific tear autoantibodies could aid in distinguishing SjD from other autoimmune and dry eye diseases. Small scale studies of autoantibodies in tears(12, 13) and saliva(14) have shown that SSA and SSB antibodies are detected in these biofluids in some seronegative patients, suggesting that tear and salivary SSA and SSB autoantibodies may be more sensitive than serum measurements. Further, symptoms of dry eye and dry mouth correlate highly with markers of inflammation in the tear fluid and saliva, respectively(15).

To understand the similarities and differences between serum and tear autoantibody expression during disease development in SjD, we analyzed tears and serum from two related murine models relative to healthy control BALB/c mice. The male non-obese diabetic (NOD) mouse spontaneously develops SjD-like AD(16) and has served as a source for identification of tear biomarkers that have been validated in human subjects(17–19). The NOD strain is reported to express serum autoantibodies to SSA and SSB(20). To account for potential confounding factors from concurrent diabetes development that occur late in AD development in this strain, we also used the haplotype-matched NOD-derived male non-obese diabetes resistant NOR/Ltj mouse(21) to investigate autoantibody composition of tear fluid and serum. The NOR mouse is a recombinant congenic strain in which limited regions of the NOD/LtJ genome are replaced by the genome of the C57Bl/KsJ strain by genetic contamination, resulting in maintenance of SjD-like AD but loss of the late-developing diabetes phenotype(21). We have previously characterized SjD-like AD development in both male NOD and NOR mice, which showed marked similarities(22). Both NOD and NOR strains exhibit sexually dimorphic disease development, where males develop earlier and more severe AD than females, and females develop earlier and more severe autoimmune sialadenitis than males(22–25). Since our primary focus is in SjD-associated AD, we thus used male mice in this study which more accurately recapitulates this disease phenotype. Our analysis identified a few common autoantibodies in both tears and serum, but, strikingly, found a discrete spectrum of autoantibodies elevated in tears versus serum in these models of SjD.

## Methods

### Animals

Three autoantibody microarrays were performed between 2020 and 2022. Both male NOD (#001976, Jackson Laboratory, Sacramento, CA) and male NOR (#002050, Jackson Laboratory, Sacramento, CA) mice were utilized as models of AD in SjD, with healthy male BALB/c (#000651, Jackson Laboratory, Sacramento, CA as controls. Consistent with previous work(16, 22, 24, 26), NOD/NOR mice aged 8- 12 weeks were considered to have early disease, mice aged 12-20 weeks to have intermediate disease, and mice aged >20 weeks to have advanced disease(26). In *Study 1,* serum was collected from male NOD and BALB/c mice with early (n=3/strain) and advanced (n=3/strain) disease. As these studies showed no major differences in serum autoantibody development across this broad age range in NOD mice, we used mice with intermediate disease for subsequent studies to minimize both minor variability in time of disease onset and comorbidities occurring later in development(27). For *Study 2*, tears and serum were collected from male NOR mice with intermediate disease (n=6) and healthy control male BALB/c mice (n=3). While serum autoantibodies were subjected to a complete analysis, tears were pooled from one, two or three mice to assess the limits of detection. For *Study 3*, tears and serum were collected from male NOD (n=5) and male NOR (n=4) mice with intermediate disease and age-matched male BALB/c mice (n=6). All study procedures conducted on animals were pre-approved by the University of Southern California’s Institutional Animal Care and Use Committee (IACUC) and were conducted in accordance with the GUIDE for Care and Use of Laboratory Animals (8^th^ Edition(28)). ARRIVE checklist is included in Supplemental Materials.

### Tissue & Biofluid Collection

Mice were anesthetized with Ketamine (60-70 mg/kg)/Xylazine (5-10 mg/kg). Tears were collected using capillaries as described previously(29) following topical carbachol administration to the LG and flash frozen. 0.5-1 mL of blood was collected by cardiac puncture using a 1 mL syringe and added to MiniCollect 0.8 mL gold cap Serum Separator tubes (Greiner Bio-One, Kremsmünster, Austria). After centrifuging the blood at 4°C, 1500 x g for 15 min, serum was obtained and stored at -80°C. For tissue collection, anesthetized mice were euthanized by cervical dislocation and LG were collected and fixed as described below.

### Immunofluorescence and Confocal Microscopy

For immunofluorescence labeling, LG were collected from male NOR and NOD mice with intermediate disease. LG were fixed, frozen in OCT, sectioned and prepared for immunolabelling as described(19). Sections were incubated overnight at 4°C with the following primary antibodies: anti- CD3e (T-cell marker, Hamster Anti-Mouse CD3e 500A2, catalog #553238, BD Biosciences, Franklin Lakes, NJ), anti-IgD (follicular B cell marker, Alexa Fluor® 647 anti-mouse IgD antibody, 11-26c.2a, catalog #405708, Biolegend, San Diego, CA), anti-B220 (B cell marker, rabbit anti-mouse RA3-6B2, catalog #MA5-48137, Thermofisher Scientific, Waltham MA), anti-CD21/CD35 (follicular dendritic cell marker, catalog# 13-0211-82, Thermofisher Scientific) and anti-PNAd carbohydrate epitope (Glycam-1 marker, catalog# 553863, BD Biosciences Franklin Lakes, NJ). Sections without primary antibodies served as negative controls. Slides were washed 3x with 0.5% Tween 20 in PBS and incubated with compatible secondary antibodies and additional probes: Alexa Fluor® 568 goat anti-hamster IgG (1:1000, catalog #A-21112), Alexa Fluor® 488 goat anti-rabbit (1:200, catalog# A- 11008), Alexa Fluor® Plus 488 donkey anti-rat (1:200, catalog# A48269TR); Alexa Fluor™ Plus 647 Phalloidin (1:200, Catalog # A30107), and DAPI (catalog #D1306), all from Thermofisher Scientific, at 37°C for 1 h then washed 3x with 0.5% Tween 20 in PBS (Catalogue# 18912014, Thermofisher Scientific). Slides were mounted with ProLong anti-fade mounting medium (Catalog # P36934, Thermofisher Scientific). Images were acquired using either a ZEISS LSM 800 confocal microscope equipped with an Airyscan detector (Zeiss, Thornwood, NY) or a ZEISS Axioscan 7 Microscope Slide Scanner and analyzed using open source Qupath software version 0.5.1. and imageJ 1.54g.

### Protein Array Profiling Analysis

Autoantigen microarrays were manufactured in the Microarray & Immune Phenotyping core Facility at the University of Texas Southwestern Medical Center (Dallas, TX, USA) and were processed as described previously(30, 31). A selection of autoantigens was made based on published literature of prior known autoantibodies(32, 33). Positive controls were also imprinted on the arrays. Each slide contained 16 identical arrays, enabling processing of 15 samples and PBS as a negative control. Arrays contained 80-120 autoantigens as well as internal controls. Mouse serum or tear samples were treated with DNAse I to remove free-DNA and then applied to autoantigen arrays at a 1:50 dilution. After incubation, autoantibody binding to the antigens on the array was detected at 532 nm (with Cy3 labeled anti-IgG) and, when IgA was analyzed, at 635 nm (with Cy5 labeled anti-IgA) to generate .tiff images. Genepix Pro 7.0 software was used to analyze the images and generate genepix report (GPR) files (Molecular Devices, Sunnyvale, California, USA). The net fluorescent intensity (NFI) of each antigen was generated by subtracting the local background and negative control (PBS) signal. The signal-to-noise ratio was also generated for each antigen. SNR is used as a quantitative measure of the ability to resolve true signal from background noise. Autoantibodies with SNR values of <3 in more than 90% of the samples were considered negative and excluded from further analysis. NFI was normalized with robust linear model using positive controls with different dilutions. NFI and SNR values as well as the R code used in analyzing the data are described in **Supplemental Methods 1**.

### Statistical Analysis

After filtering autoantibodies with SNR <3 for a given mouse group in each array, remaining NFIs were further normalized by calculating row trimeans and modeling the relationship with the residuals by fitting a quantile regression by using the Limma(34) voom package for *Studies 2 and 3* or the code provided in the **Supplemental Methods** based on Kiripolsky et. al.(35) *for Study 1,* depending upon the distribution of the raw data. Normalized values were log transformed and scaled using the biweight midvariance method to account for variances in scaling. Differential autoantigen expression was determined using the Wilcoxon Rank Sum test and correction for multiple comparisons was done using the Benjamini and Hochberg procedure. The Spearman correlation coefficient was calculated for IgG vs IgA in mouse tears in *Study 3*.

## Results

### TLS in LG of NOD and NOR

TLS can form within target organs undergoing chronic inflammatory/autoimmune processes, whose primary function is not lymphogenic in nature. Like germinal centers (GC), they are lymphoid structures where memory lymphocytes and/or precursors can be re-stimulated by antigen which results in clonal expansion(36). We have also shown that miR-155-5p, a marker of GC development(37), is significantly upregulated in the LG of male NOD mice, particularly in the infiltrating lymphocytes of the NOD LG(38). In this report, we evaluated the presence of TLS in LG of two murine models of AD in SjD, male NOD and NOR mice. **Fig. 1** shows representative images from LG from mice of each strain with intermediate disease. Labeling of lymphocytic infiltrates with IgD, a marker of naive follicular B-cells(39, 40) (**Fig. 1A),** and B220, a general marker for B cells(41) (**Fig. 1B**), showed enriched B cell zones enveloping CD3e positive T cells (**Fig. 1A, B**) in the LG. Some foci showed a clear segregation of T and B cells in a GC-like manner. These infiltrates were also enriched in other components of TLS including markers of high endothelial venules such as PNAd(42) (**Fig 1C)** and highly compartmentalized CD21/CD35 positive follicular dendritic cells (**Fig. 1D**). TLS were detected in LG from both strains at early, intermediate, and advanced disease, increasing gradually with age (data not shown).

**Figure 1.**
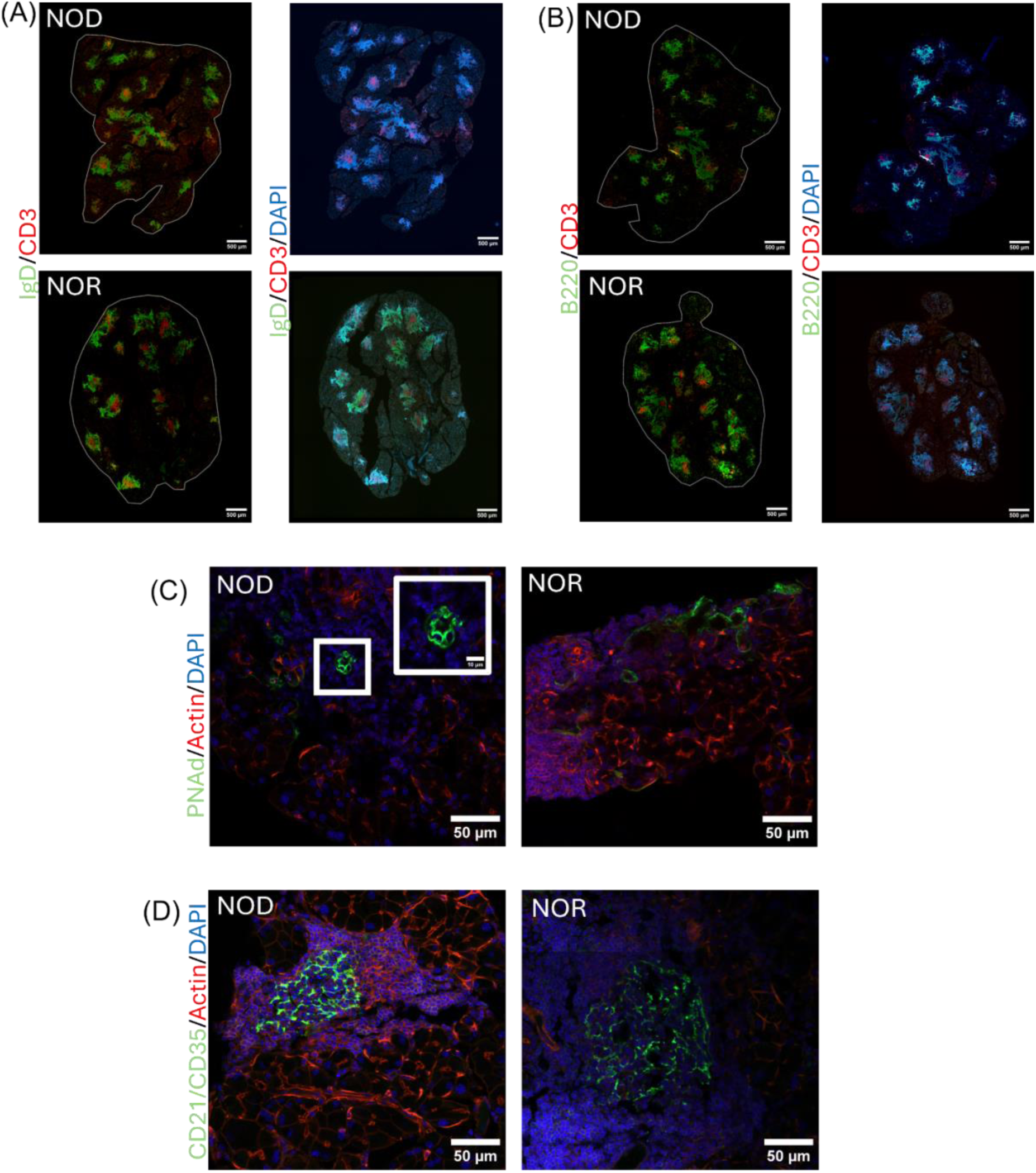
Representative images of tertiary lymphoid structures (TLS) in LG of male NOD and NOR mice with intermediate disease. Immunofluorescence (IF) images show components of TLS as follows: **(A)** Antibodies to IgD highlighting follicular B cells (green) and CD3 highlighting T cells (red). DAPI labels nuclei (blue). **(B)** Antibodies to B220 highlighting B cells, (green) and CD3 highlighting T cells (red). DAPI labels nuclei (blue). **S**cale bar in **(A)** and **(B)**, 500 µm. (C) Antibody to PNAd, highlighting high endothelial venules (green) and Alexa Fluor™ Plus 647 Phalloidin highlighting F-actin (red). DAPI labels nuclei (blue). **(D)** Antibodies to CD21/CD35, highlighting follicular dendritic cells green) and Alexa Fluor™ Plus 647 Phalloidin highlighting F-actin (red). DAPI labels nuclei (blue). Scale bars in **(C)** and **(D)**, 50 µm.

### Autoantibodies in Serum of NOD and NOR mice

Three separate arrays were conducted to compare IgG autoantibody abundance in both tears and serum from male NOD and NOR mice versus healthy control BALB/c mice. A complete list of autoantigens, their abbreviations, and disease association is provided in the **Supplemental Data 1** file. *Study 1* compared serum autoantibody abundance from mice with early and advanced disease in male NOD mice versus age-matched healthy control BALB/c mice. NOD mice at both disease states exhibited significantly increased serum IgG reactivity against four common autoantigens relative to age-matched healthy male BALB/c mice **(Supplemental Table 1)**. These included – Polymyositis/Scleroderma antigen 100 (Pm/Scl-100), Proliferating Cell Nuclear Antigen (PCNA), Anti-liver cytosolic antigen type 1 (LC-1), and Heterodimer Ku protein subunits 70 & 80 (KU P70/P80). Additionally, Small RNA Binding Exonuclease Protection Factor La (La/SSB), threonyl-tRNA synthetase (PL7), Thyroid Peroxidase (TPO), Intrinsic Factor (IF) and Mitochondrial Antigen had significantly elevated serum IgG reactivity at early disease, with greater variability in advanced disease **(Supplemental Table 1)**. For each of the nine antigens above, NOD mice with early and advanced disease showed the same significant differences between healthy (BALB/c) and diseased (NOD) mice (**Supplemental Table 1**). Therefore, age was not considered a confounder for development of these autoantibodies. The common elevation of these autoantibodies across the spectrum of SjD-associated AD development suggests that they form early in disease and remain elevated. These findings further suggested that use of mice with intermediate disease might capture changes characteristic of AD in SjD, but avoid confounders associated with advanced disease and development of comorbidities occurring >20 weeks of age(27).

Other autoantibodies were elevated both in the serum of advanced disease NOD mice and age-matched BALB/c mice, namely Signal Recognition Particle 54 kDa protein (SRP54) and Major centromere autoantigen B (CENP-B), (**Supplemental Fig. 1**). This suggests their formation is dependent on age rather than on disease progression. Additionally, a slight decrease was observed in PM/Scl-100 and Glycated Albumin autoantibodies with age (**Supplemental Fig. 1**).

In *Study 2,* we used NOR mice with intermediate disease. We identified increased immunoreactivity in NOR mouse serum relative to serum of healthy age-matched male BALB/c mice against sixteen autoantigens (**Supplemental Table 2**). **Fig. 2A** shows the combined results of the analyses from *Studies 1 and 2*, which highlights elevations in serum autoantibodies common to both NOD and NOR mice. Common IgG autoantibodies distinct to serum included PM/Scl 100, LC1, La/SSB, Mitochondrial Antigen and PCNA, relative to serum from healthy BALB/c controls. In *Study 3*, we reconfirmed IgG and added IgA immunoreactivity against 80 autoantigens in serum from male NOD and NOR mice with intermediate disease relative to healthy BALB/c controls. This study validated significantly increased serum IgG reactivity against LC1, PCNA and La/SSB in both male NOD and NOR mice (**Fig. 2B**) compared to BALB/c. Thus, these serum autoantibodies were common to NOD and NOR mouse serum across all studies. In addition, arrays used in *Study 3* had antigens not present in previous studies including several exclusive to serum (**Supplemental Figure 2**).

**Figure 2.**
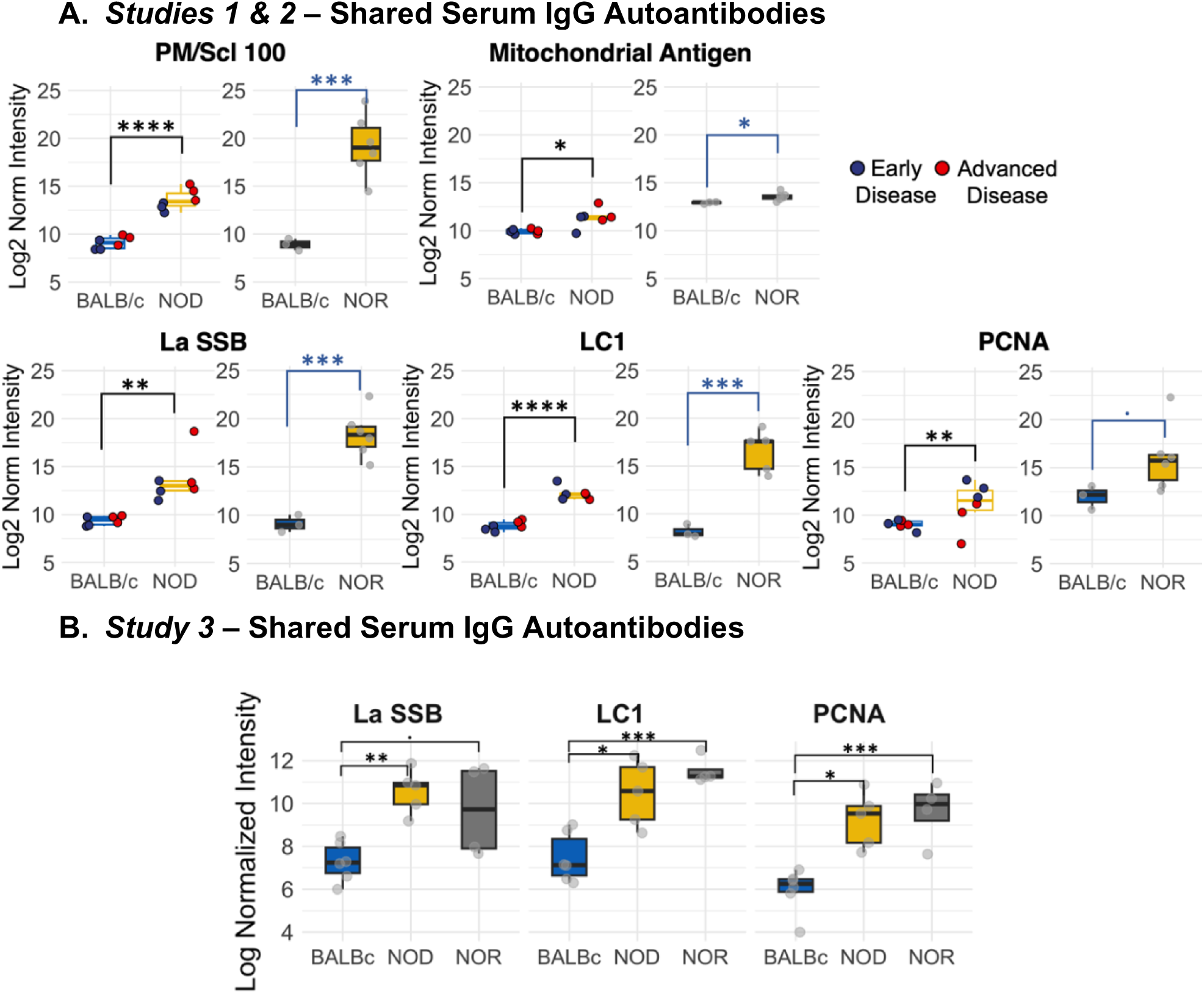
Differential IgG autoantibody expression of autoantibodies in serum in male NOD and NOR mice. **(A)** Box plots showing autoantibodies significantly increased in serum of male NOD (N=6) mice as compared to male BALB/c (N=6) mice from mice in two different age groups from *Study 1* (left graphs). Box plot showing log normalized intensities of autoantibodies significantly upregulated in serum of male NOR mice with intermediate disease (N=6) as compared to age-matched male BALB/c mice (N=3) from *Study 2* (right). **(B)** Box plots showing validated serum IgG autoantibodies increased in serum only from male NOD (n=5) and male NOR (n=4) mice with intermediate disease and age-matched male BALB/c mice (n=6) from *Study 3.* Moderated t-statistics estimated using Limma R package. (* p_adj_ < 0.05, ** p_adj_ < 0.01, *** p_adj_ < 0.001, **** p_adj_ < 0.0001). *Abbreviations: LC1 – Anti-liver cytosolic antigen type 1; PM/Scl-100 – polymyositis/ Scleroderma antigen 100; PCNA – Proliferating Cell Nuclear Antigen; La/SSB – Small RNA Binding Exonuclease Protection Factor La*

Compared to total serum IgG, total serum IgA was detected at a significantly lower level in male NOD and NOR mice (p < 10^-4^, One-way ANOVA, **Supplemental Figure 3A**). The only antigens with significantly elevated IgA reactivity in serum of both NOD and NOR mice with intermediate disease were intrinsic factor (IF) and single stranded DNA (ssDNA) (**Supplemental Fig. 3B**). A modest increase was also observed for Cytochrome C (Cyt C), Gliadin and Glycoprotein 2 (GP2) (**Supplemental Table 3**).

### Autoantibodies in Tears and Serum of NOD and NOR mice

*Studies 2* and *3* included tear samples in addition to serum. Several IgG autoantibodies were upregulated in both serum and tears, including autoantibodies to PL7 and KU (P70/P80) in *Study 2* (**Supplemental Fig. 4A)**, and PL7, Glutamic Acid Decarboxylase 65 (GAD65), and Liver kidney microsome type 1 (LKM1) in *Study 3* (**Supplemental Fig. 4B)**. The only common autoantibody seen in both *Studies 2 and 3* in serum and tears was to PL7. Additional serum autoantibodies, including against intrinsic factor (IF) may be biologically relevant as they were increased across multiple assays and different antibody isotypes but not significant in *Study 2* (**Supplemental Table 2**).

### Autoantibodies in Tears of NOD and NOR mice

We explored the tear autoantibody profile of IgG and IgA isotypes in tears of both male NOD and NOR mice with intermediate disease versus tears of healthy control BALB/c mice **(Table 1).** In *Study 3*, we found 6 IgG autoantibodies that were significantly upregulated in tears of both NOD and NOR mice. These include autoantibodies to Histidyl-tRNA synthetase (Jo-1), Anti-Islet Cell Antigen 512 (IA-2), Tissue Transglutaminase 2 (tTG), Dermatomyositis specific autoantigen Mi-2 (Mi-2), thyroid peroxidase (TPO) and SAE1/SAE2 (**Fig. 3A**).

**Figure 3.**
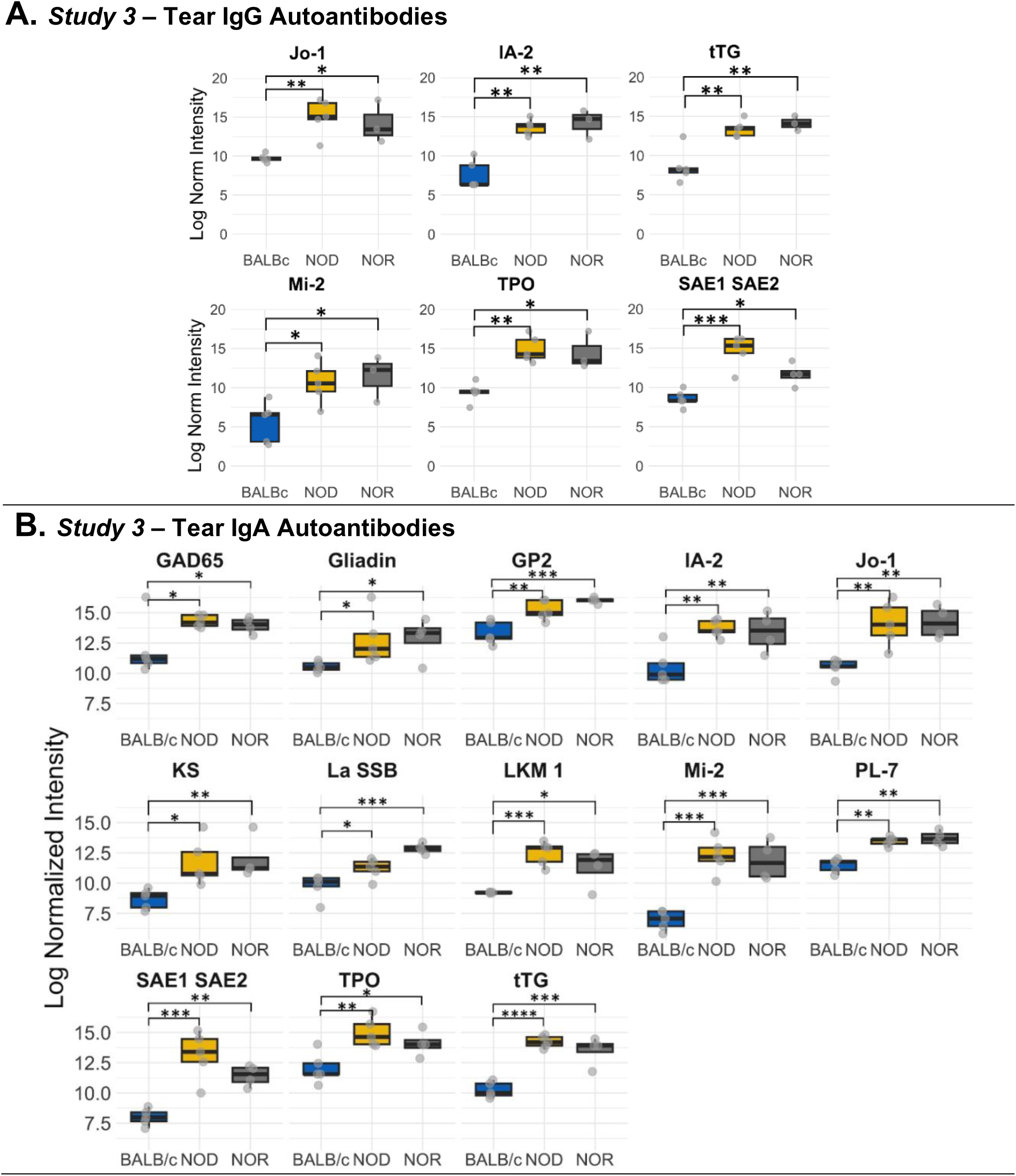
Differential autoantibody expression of autoantibodies unique to tears of male NOD and NOR mice. **(A)** IgG autoantibodies that are unique to tears from male NOD (N=5) and male NOR (N=3) mice with intermediate disease versus male BALB/c (N=3) mice from *Study 3*. Moderated t-statistics estimated using Limma R package. **(B)** Differentially expressed IgA autoantibodies in tears of the same male NOD, male NOR and BALB/c mice from *Study 3*. Moderated t-statistics estimated using Limma R package (ref). p-values adjusted by Benjamini and Hochberg procedure to account for multiple comparisons (* p_adj_ < 0.05, ** p_adj_ < 0.01, *** p_adj_ < 0.001, **** p_adj_ < 0.0001). *Abbreviations: Jo-1 – Histidyl-tRNA synthetase; IA-2 – protein tyrosine phosphatase-like Anti-Islet Cell Antigen 512; tTG – Tissue Transglutaminase 2; GAD65 – Glutamic Acid Decarboxylase 65; LKM1 – Liver kidney microsome type 1; Mi-2 – Dermatomyositis specific autoantigen Mi- 2; PL-7 – threonyl-tRNA synthetase; TPO – Thyroid peroxidase; GP2 – Glycoprotein 2; IF – Intrinsic Factor; KS – anti-asparaginyl-tRNA synthetase or KS; La/SSB – Small RNA Binding Exonuclease Protection Factor La; SAE1/SAE2 – SUMO1 activating enzyme subunit 1/2*

**Table 1.** Differentially Expressed IgG & IgA Autoantibodies in Tears from Mice in *Study 3*.

|  | NOD vs BALB/c |  |  |  | NOR vs BALB/c |  |  |  |
| --- | --- | --- | --- | --- | --- | --- | --- | --- |
|  | IgG |  | IgA |  | IgG |  | IgA |  |
|  | p<sub>adj</sub> | Log<sub>2</sub> FC | p<sub>adj</sub> | Log<sub>2</sub> FC | p<sub>adj</sub> | Log<sub>2</sub> FC | p<sub>adj</sub> | Log<sub>2</sub> FC |
| tTG | **0.0035** | 4.23 (18.76) | **3.01E-05** | 6.69 (103.25) | **0.0011** | 3.89 (14.82) | **3.30E-04** | 5.48 (44.63) |
| Mi-2 | **0.0164** | 3.12 (8.70) | **1.01E-04** | 5.80 (55.71) | **0.0118** | 3.23 (9.38) | **3.63E-04** | 5.17 (36.00) |
| Jo-1 | **0.0038** | 4.24 (18.90) | **0.0002** | 4.44 (21.70) | **0.0139** | 3.60 (12.12) | **0.0019** | 4.44 (21.70) |
| IA-2 | **0.0018** | 4.62 (24.60) | **0.0004** | 4.21 (18.50) | **0.0047** | 3.93 (15.24) | **0.0042** | 3.90 (14.93) |
| GP2 | 0.1438 | 1.80 (3.48) | **0.0008** | 3.87 (14.62) | **0.0081** | 3.91 (15.03) | **5.96E-04** | 5.17 (36.00) |
| SAE1 SAE2 | **4.87E-04** | 5.36 (41.06) | **1.51E-04** | 5.80 (55.71) | **0.0421** | 2.35 (5.10) | **0.0056** | 3.40 (10.56) |
| IF | 0.2126 | 1.34 (2.53) | 0.0942 | 2.02 (4.05) | **0.0078** | 3.71 (13.08) | **0.0010** | 4.47 (22.16) |
| TPO | **0.0041** | 4.27 (19.30) | **0.0021** | 3.47 (11.08) | **0.0335** | 3.04 (8.22) | **0.0190** | 2.61 (6.10) |
| PL-7<sup>†</sup> | **0.0129** | 2.73 (6.63) | **0.0030** | 3.08 (8.45) | 0.0640 | 2.10 (4.29) | **0.0015** | 3.55 (11.71) |
| KS | 0.0973 | 1.80 (3.48) | **0.0126** | 3.23 (9.38) | **0.0126** | 2.85 (7.21) | **0.0098** | 3.51 (11.39) |
| GAD65<sup>†</sup> | **0.0123** | 3.18 (9.06) | **0.0149** | 3.01 (8.05) | 0.0816 | 2.06 (4.17) | **0.0434** | 2.49 (5.62) |
| GP210 | **0.0292** | 2.28 (4.85) | 0.5132 | 1.04 (2.05) | **0.0041** | 5.86 (50.67) | **0.0105** | 3.46 (11.00) |
| LKM 1<sup>†</sup> | 0.1438 | 2.20 (4.60) | **0.0049** | 4.68 (25.63) | 0.2937 | 1.68 (3.20) | **0.0176** | 2.90 (7.46) |
| gDNA | 0.0777 | 2.78 (6.87) | 0.2038 | 2.69 (6.45) | **0.0257** | 3.20 (9.19) | 0.0579 | 2.87 (7.31) |
| Gliadin | 0.0973 | 2.28 (4.85) | **0.0461** | 2.75 (6.73) | 0.1056 | 2.20 (4.60) | **0.0500** | 2.62 (6.15) |
| La SSB | 0.8167 | -0.23 | **0.0419** | 1.86 (3.63) | **0.0201** | 2.09 (4.26) | **5.28E-04** | 4.75 (26.91) |
| NXP2 | 0.5498 | 0.61 (1.52) | 0.5203 | 0.82 (1.76) | **0.0002** | 4.08 (16.91) | **0.0056** | 3.48 (11.16) |
| PM Scl75 | **0.0189** | 3.16 (8.94) | 0.0615 | 2.72 (6.59) | **0.0362** | 2.47 (5.54) | 0.4236 | 0.81 (1.75) |
| T1F1g | **0.0403** | 3.60 (12.50) | - | - | **0.0395** | 3.83 (15.68) | - | - |
| LC1 | - | - | 0.1674 | 1.90 (3.73) | - | - | **0.0280** | 2.36 (5.13) |
| dsDNA | - | - | 0.0979 | 2.27 (4.82) | - | - | **0.0108** | 2.34 (5.06) |
| TNF a | - | - | **0.0063** | 3.03 (8.17) | - | - | 0.1389 | 1.54 (2.90) |
| Cardiolipin | 0.0855 | 2.14 (4.41) | - | - | **0.0168** | 3.28 (9.71) | - | - |
| SmD1 | **0.0024** | 3.74 (13.36) | - | - | **0.0301** | 2.33 (5.02) | - | - |

*p<sub>adj</sub> < 0.05 was considered significant, multiple comparisons p-values were adjusted by the Benjamini-Hochberg procedure.*
*Statistically significant p-values are shown in bold.*
<sup>†</sup>Also upregulated in serum of male NOD and NOR mice compared to healthy BALB/c

In *Study 3,* IgA reactivity was tested against the same panel of 80 autoantigens. There was significantly higher IgA reactivity against 14 autoantigens in both male NOD and NOR mouse tears relative to healthy BALB/c tears, of which 13 were unique to tears. These included autoantibodies to tTG, Mi-2, Jo-1, IA-2, GP2, SAE1/SAE2, TPO, PL-7, KS, GAD65, LKM1, La/SSB and Gliadin (**Fig. 3B**). While there was largely an agreement between the upregulated IgG and IgA autoantibody species (**Table 1**), tears of male NOR mice had significantly higher total IgA than total IgG (p=0.002, two-way ANOVA), with comparable levels of IgA and IgG in the healthy control male BALB/c (**Supplemental Fig. 5A**). Reactivity to IgA control was significantly higher than reactivity to IgG controls in tears of all mice but was relatively higher in the male NOD and NOR mice compared to BALB/c (**Supplemental Fig. 5B)**. Positive correlations between the signal intensities for IgGs and IgAs to the same autoantigens were observed for 70 out of 80 autoantigens tested (**Supplemental Methods 1**) 50 of which were statistically significant. The highest positive correlations were observed for IgGs and IgAs to the six shared epitopes in NOD and NOR, but not for BALB/c controls (**Supplemental Figure 6**).

## Discussion

Serum expression of SSA and SSB autoantibodies is observed in ∼25-75% of SjD patients(43, 44), respectively, while these autoantibodies are also detected in patients with systemic lupus erythematosus and rheumatoid arthritis(9, 45). This lack of sensitivity and specificity is a continual challenge for SjD diagnosis. The possible identification of additional diagnostic autoantibodies in SjD has been previously investigated, with researchers reporting serum autoantibodies to SP1, Ca6 or PSP with significantly better sensitivity than antibodies to Ro or La(46). In this study we show for the first time that two related murine disease models of SjD exhibiting AD, the male NOD and NOR mice, express autoantibodies that are largely distinct in serum versus tears. The autoantibodies unique to serum were primarily IgG autoantibodies. However, the autoantibodies unique to tears included both IgG as well as IgA isotypes, while the autoantigens recognized by the IgG tear autoantibodies were all represented in the tear IgA autoantibodies.

IgA is the second most common antibody isotype in serum, after IgG(47). Nearly 90% of the IgA found in serum is monomeric IgA1, produced in the bone marrow, which plays no role at the mucosal surface. However, the IgA found in mucosal tissues, such as the LG is secreted dimeric IgA (dIgA), produced by resident plasma cells in the glandular interstitium. dIgA binds to polymeric Immunoglobulin Receptor (pIgR) expressed on the basolateral membrane of polarized LG acinar cells and is transcytosed to the apical surface where pIgR is cleaved and dIgA released into tears as secretory IgA (sIgA)(48). In SjD, TLS are found in LG(49). IgA autoantibodies generated by solitary plasma cells at early disease stages as well as from local TLS in established disease would therefore be recovered in tears due to this robust active transport process.

IgG present in tears is either produced in the LG or originates from the circulation. IgG is not known to be actively transported, and its presence in tears is likely due to passive paracellular leakage. Such leakage may increase with disease progression due to degradation of extracellular matrix, epithelial cell apoptosis, and diminishing tissue organization(49). The plausibility of detecting SjD autoantibodies in tears was previously shown in a small study by Toker et. al(12) that demonstrated that roughly 16 of the 28 patients positive for serum SSA and 14 of the patients positive for serum SSB were positive for SSA and SSB respectively, in tears. However, this study only investigated IgG autoantibodies. The Toker study and similar studies conducted in saliva have also shown the presence of SSA and SSB in tears or saliva of seronegative patients(12–14). To our knowledge, no investigations have studied the complete spectrum of autoantibodies present in tears, nor compared them to the spectrum of serum autoantibodies in SjD patients or SjD disease models. We found that the autoantibody repertoire of male NOD and NOR mice tears was quite distinct from that found in their serum. Increased IgG and IgA reactivity to Jo1, IA2, tTG, Mi-2, TPO and SAE1/SAE2 was seen only in tears, but not in serum.

In the healthy eye, conjunctiva associated lymphoid tissue (CALT) comprises a secondary lymphoid organ (SLO) that is crucial to ocular defense against pathogens. CALT has been studied in healthy BALB/c and C57/BL6 mice with changes observed in their number and size upon exposure to pollen(50), induction of evaporative dry eye disease(50) or with age(51).

We suspect that autoantibodies observed in the tears of male BALB/c mice in this study may be produced by CALT in addition to resident LG plasma cells to maintain ocular health. On the other hand, TLS, while having features similar to SLO, develop in organ-specific autoimmune diseases such as the SG and LG of patients with SjD(36, 52) where they are not normally present. Their formation involves B-cell activation and differentiation into autoantibody producing plasma cells via a series of somatic hypermutation and class switch recombination of the Ig genes(1). These structures actively support molecular machinery responsible for pathogenic autoantibody production. Our data show the presence of characteristic TLS within LG of male NOR and NOD mice. We also observe expression of other components of TLS including PNAd, a marker of high endothelial venules in the dark zone(42), and CD21/CD35, a marker of follicular dendritic cells of the light zone(53). The tear-specific IgA and IgG autoantibodies seen in NOD and NOR mice may be largely produced from LG TLS, although a recent study has also suggested that CALT of patients with autoimmune eye-related diseases including SjD and graft-versus-host disease can produce tear autoantibodies to citrullinated proteins(54). In either case, tear IgA autoantibodies appear to be generated locally since they differ both in predominant isotype and antigen from those expressed in serum in the same animals.

An important question is to understand how tear autoantibodies are related to disease development and progression in SjD. Many studies have suggested that the autoantigens triggering production of SSA and SSB autoantibodies, Ro and La, are released locally in exocrine tissue due to early disease triggers (viral infection, ER stress, apoptosis) where they can be processed in TLS to generate autoantibodies(55). Other anatomical features of TLS suggest that local TLS are most likely to express local autoantigens(36). For instance, early events in antigen presentation in exocrine glands are facilitated by the epithelial cells themselves, through the process of autoimmune epithelitis(56). TLS may have poorly developed lymphatics relative to the more organized Secondary Lymphoid Organs (SLO) in lymphoid tissue. In the absence of these lymphatics, mature dendritic cells carrying antigen from other peripheral sites may not be able to reach TLS.

With this process in mind, we considered the common IgA and IgA tear autoantigens– Jo-1, IA2, tTG, Mi2, SAE1/SAE2 and TPO. Synergism of IgG and IgA autoantibodies to common autoantigens is known to potentiate interferon responses in systemic lupus erythematosus(57). Given the high degree of correlation observed between IgG and IgA for these six shared antigens in tears of SjD model mice, their role in synergistically propagating autoimmunity in LG requires further investigation. The same apoptotic processes that may lead to exposure of Ro and La may expose additional autoantigens leading to autoantibody development to Jo-1, tTG, Mi2 and SAE1/SAE2. Anti-Jo-1 IgG and IgA autoantibodies target histidyl-tRNA synthetase(58), a major biosynthetic enzyme. Serum IgG elevation of this antigen is associated with anti-synthetase syndrome(59–61). Mi2 autoantigen is a key marker of antimitochondrial autoantibodies. These antibodies have been strongly associated with primary biliary cholangitis(43). Notably, anti-Jo-1 and Mi2 IgG were found to be absent in sera of 26 SjD patients and were only detected in patients with myositis(62), Similarly, anti- SAE1/SAE2 IgG is reported to be exclusively found in serum of patients with Myositis. Tissue transglutaminase (tTG) is an abundant protein found in multiple tissues in both intracellular and extracellular spaces that catalyzes protein cross linking(63). IgA autoantibodies to tTG are commonly used to diagnose celiac disease in children(64). However, it is notable that autoantibodies to this protein are detected in saliva from SjD(65).

It is unclear how local autoantigens could contribute to tear autoantibodies to thyroid peroxidase (TPO), a tissue-restricted thyroid secretory protein important in generation of thyroglobulin or IA-2, an islet-specific autoantigen for type 1 diabetes(66, 67). Autoantibodies to TPO are commonly seen in autoimmune thyroid diseases such as Hashimoto’s disease(68). Interestingly, diabetes, autoimmune thyroiditis, myositis(69, 70) and primary biliary cholangitis (associated with autoantibody to Mi2, above) have been explored as potential co-morbidities in SjD (27, 71), Autoimmune thyroiditis is a co- morbidity in the NOD mouse in 5-15% of animals(27), while diabetes is a defining phenotype in the NOD mouse. While the presence of autoantibodies to these proteins is unsurprising given co- morbidities of the disease model, their concentration in tears instead of in serum is surprising.

Previously, studies have found salivary anti-muscarinic receptor autoantibodies had higher sensitivity in diagnosing younger patients with SjD at an earlier stage of disease progression(72). Another study validated salivary anti-histone and anti-tTG autoantibodies in 34 patients with SjD with an area under the receiver operator curve (AROC) of 0.87 or higher(65), which is significantly better than the AROC for serum SSA and SSB. Adding tTG and histone autoantibodies to salivary Ro and La gave an AROC of 0.99(65). That study also found significant elevation of anti-cardiolipin autoantibodies in SjD saliva when compared to healthy controls and SLE patients. This lends further credence to the utility of testing local biofluids in addition to systemic biofluids in SjD. Thus, we propose tear autoantibodies may outperform serum biomarkers in sensitivity and specificity for ocular manifestations of SjD. Future studies will explore additional aspects of tear autoantibody development including variability in the time course of expression associated with disease progression and also explore the presence of these hits in SjD patients.

## Author Contributions

JAM and SHA secured research funding and contributed to experimental design. YJ, SK, and ME were also involved in the study conception and design. SK, YJ, ME and SA acquired the murine samples, and IR and SA carried out the biochemical analysis of samples and acquisition of raw data. CZ and SK conducted the data analysis. SK, SA, IR, CZ, SHA, ME, and PR were involved in the interpretation of data. SK, SA, ME and SHA wrote the first draft. All authors were involved in revising the final article for important intellectual content, and all authors approved the final version to be published.

## Data Availability Statement

The datasets generated for this study can be found in the NCBI Gene Expression Omnibus (GEO). The code used for analyzing this dataset can be found in the GitHub repository at https://github.com/singhkakan/Autoantibody_Project and has also been provided as Supplementary Methods 1 file.

## Supplementary Material

Supplemental Figures and Tables. (Word document)

Supplemental Methods 1 (PDF document)

Supplemental Methods 2 (Excel document)

---

**Supplemental Figure 1.**
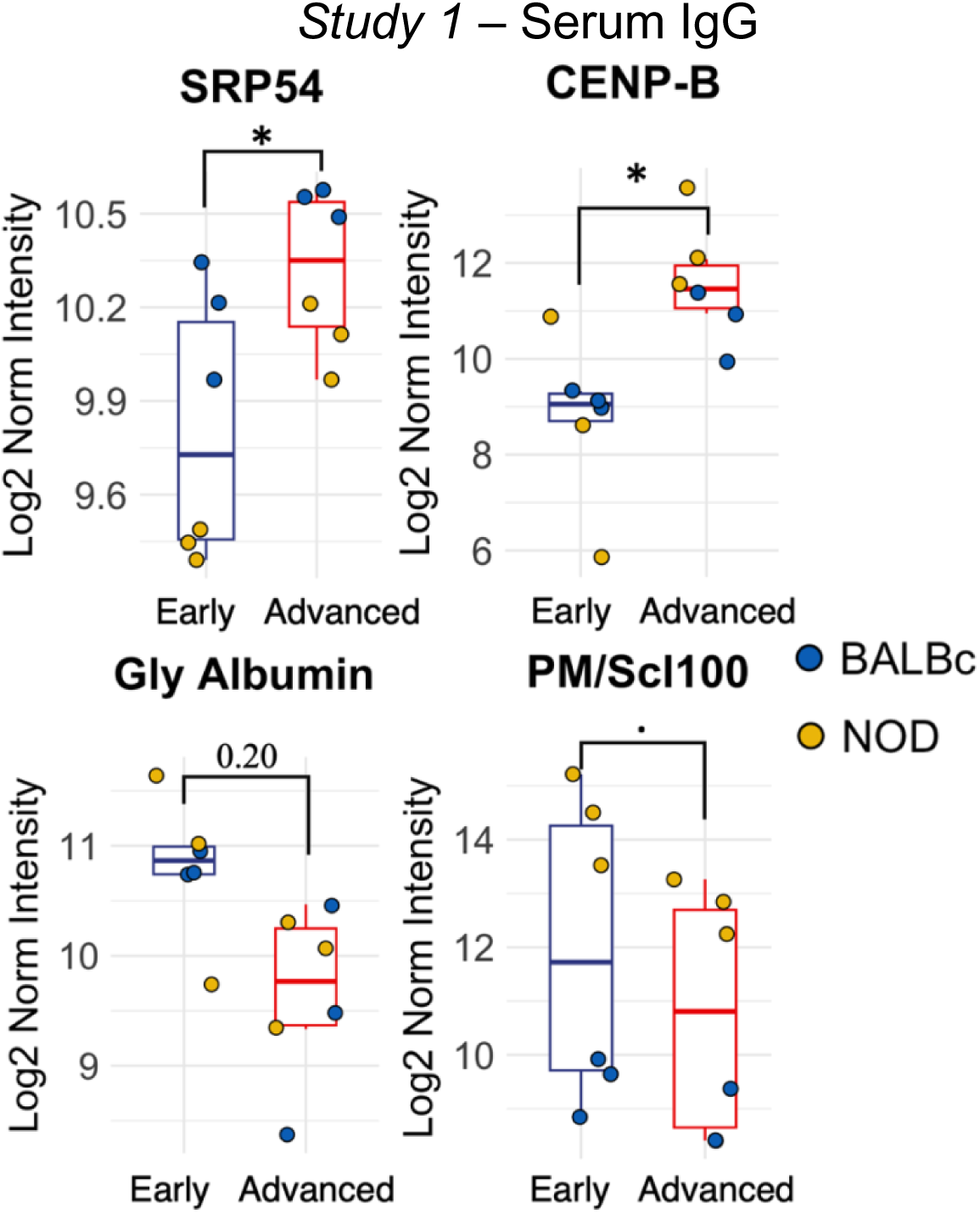
Effect of age on serum IgG autoantibodies in male NOD mice with early disease (n=3) and advanced disease (n=3) versus BALB/c mice (n=3 per age group matched to NOD mice). Each point represents one mouse. Moderated t-statistics estimated using Limma R package. (.padj < 0.1, * padj < 0.05, ** padj < 0.01, *** padj < 0.01). *SRP54 – Signal Recognition Particle 54 kDa protein; CENP-B – Major centromere autoantigen B (Centromere protein B); PM/Scl-100 – polymyositis/ Scleroderma antigen 100; Gly Albumin – Glycated Albumin;*

**Supplemental Figure 2.**
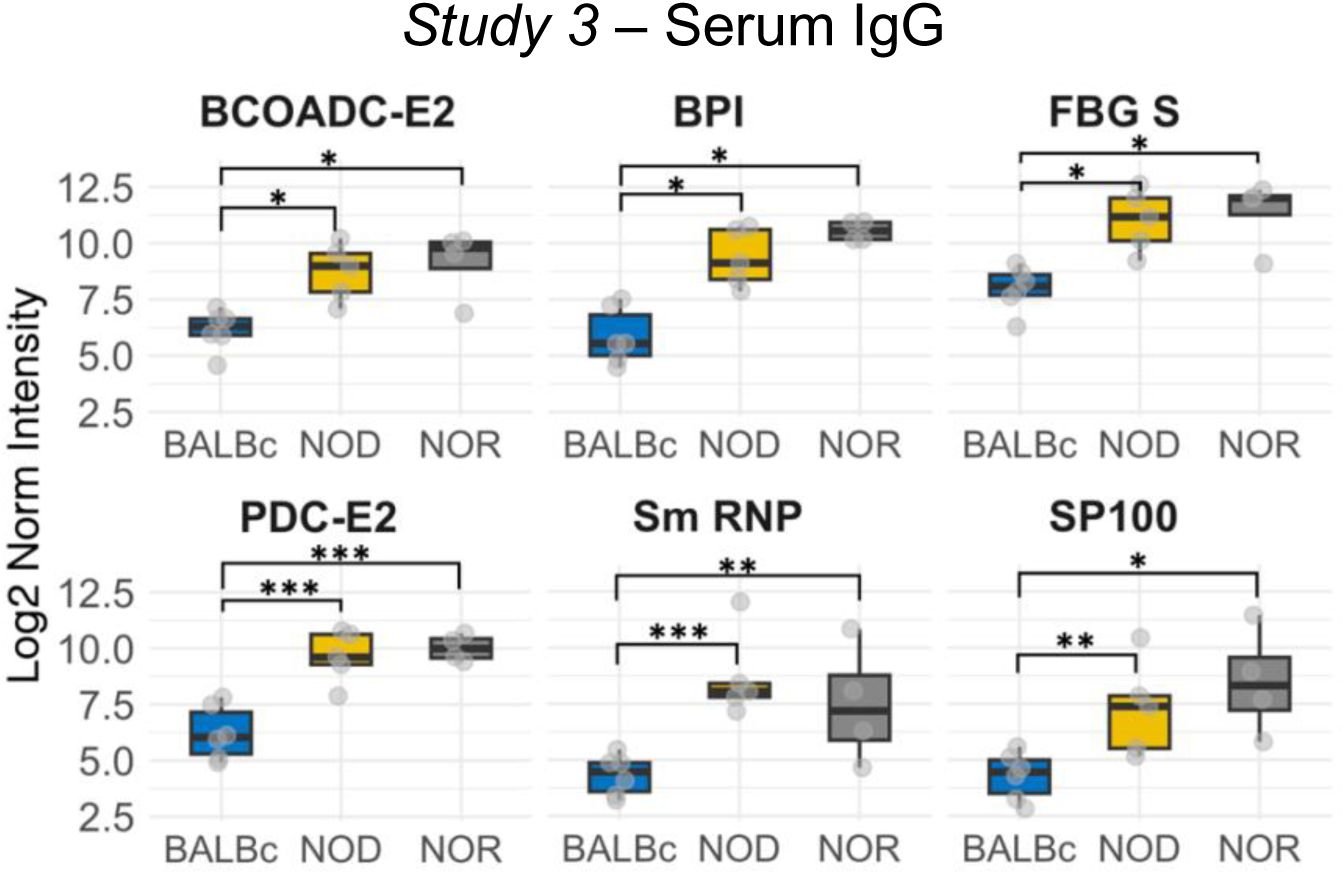
Additional significantly upregulated IgG autoantibodies from *Study 3* that were not assessed in the arrays from *Studies 1 and 2*. Each point represents one mouse with tears collected from male NOD mice (n=5) and male NOR mice (n=4) with intermediate disease versus age-matched BALB/c (n=6) mice. Moderated t-statistics estimated using Limma R package. (.padj < 0.1, * padj < 0.05, ** padj < 0.01, *** padj < 0.001). *BCOADC – Branched chain 2-oxo acid dehydrogenase complex; Bactericidal/permeability-increasing protein (BPI); FBG S – Fibrinogen Type I – S; PDC-E2 – Pyruvate dehydrogenase complex component E2; Sm RNP – Smith/Ribonuclear Protein Antibody (anti-ENA); SP 100 – Anti sp100 nuclear antigen*

**Supplemental Figure 3.**
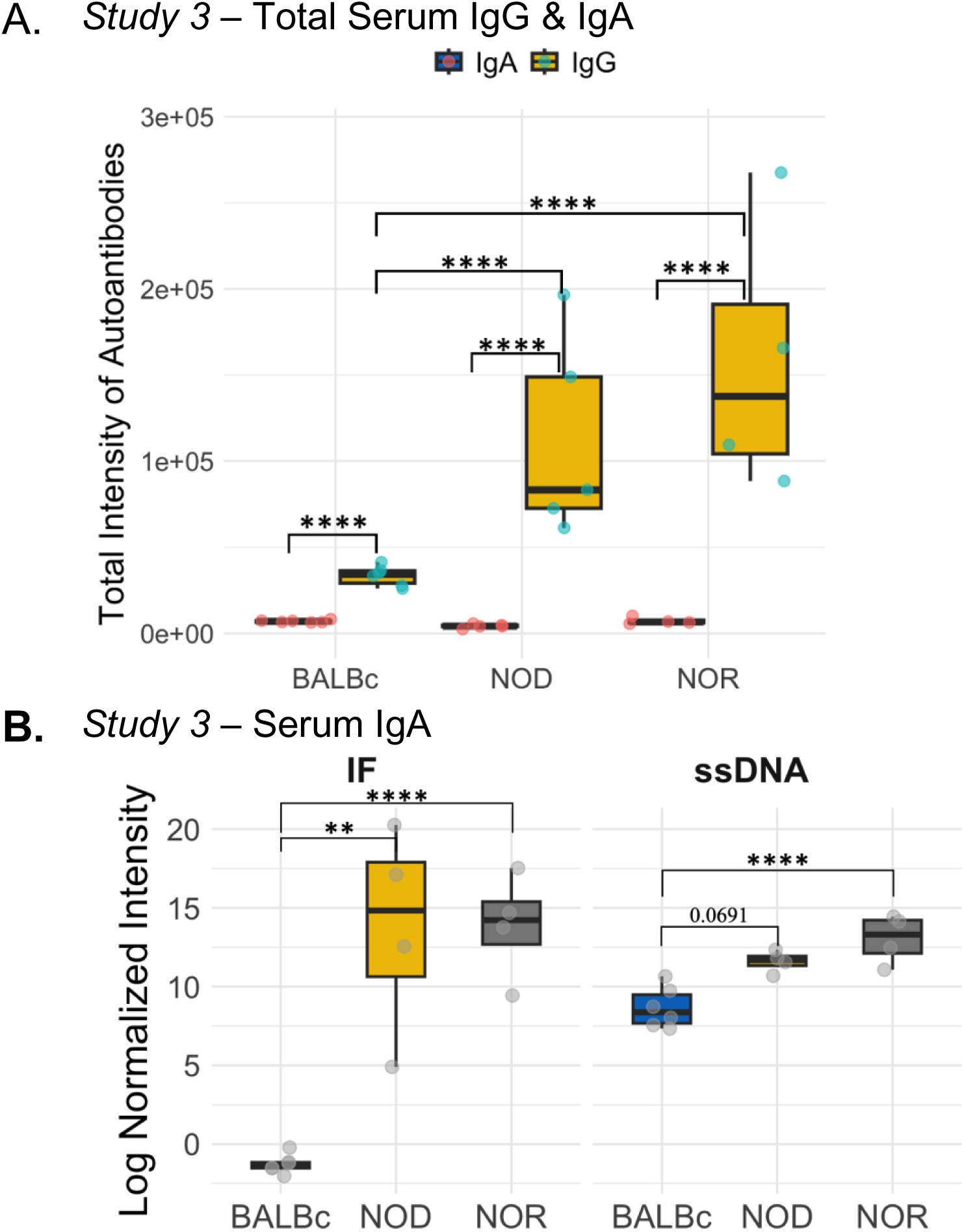
S**e**rum **IgG and IgA levels per mouse strain from *Study 3.* (A)** Boxplots showing sum of raw signal intensity of Ig reactivity for 80 autoantibodies for isotypes IgA or IgG in mouse serum from *Study 3*. (**** p_adj_ < 0.0001, One-way ANOVA with Tukey’s Honest Significant Difference Test for multiple comparison correction). **(B)** Differentially expressed IgA autoantibodies in serum of adult male NOD, male NOR and BALB/c mice from *Study 3*. Each point represents one mouse, with serum and tears collected from male NOD mice (n=5) and male NOR (n=4) with intermediate disease versus age-matched male BALB/c (n=6) mice. Moderated t-statistics estimated using Limma R package. Adjusted p-values calculated using the Benjamin & Hochberg Procedure with alpha = 0.05 (* p < 0.05, ** p < 0.01, *** p < 0.001). *Abbreviations: IF – Intrinsic Factor; ssDNA – Single stranded DNA;*

**Supplemental Figure 4.**
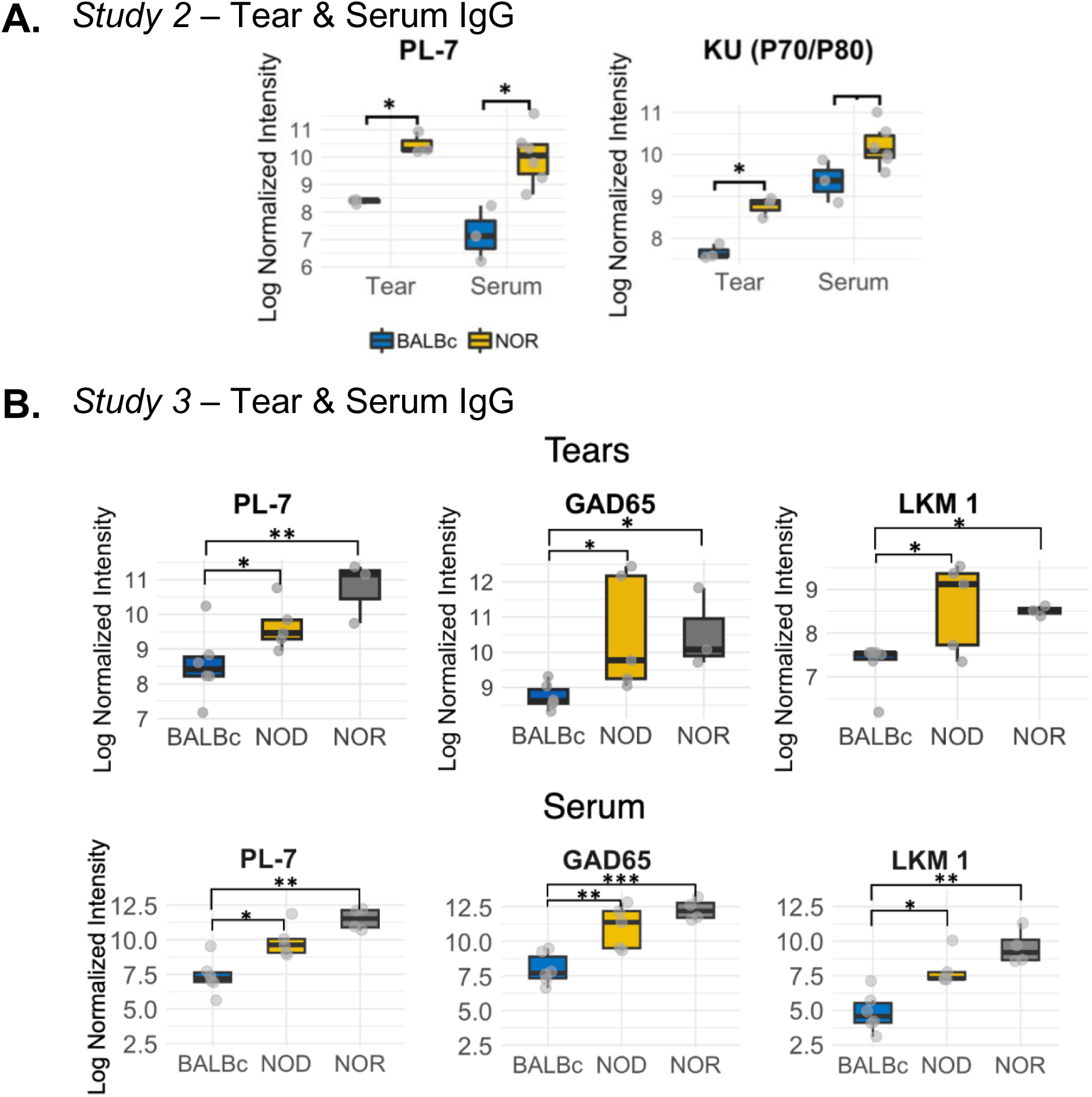
IgG Autoantibodies common to serum & tears. **(A)** IgG Autoantibodies upregulated in serum and tears of male NOR mice with intermediate disease relative to age-matched male BALB/c mice from *Study 2*. For serum collection, each point represents one mouse with serum collected from NOR (n=6) and BALB/c (n=3) mice. Tears were pooled from one, two or three NOR mice per sample (n=3 samples), with tears collected from one mouse per sample for BALB/c controls (n=3) **(B)** IgG Autoantibodies upregulated in tears and serum of male NOD and NOR mice from *Study 3.* Each point represents one mouse, with serum and tears collected from male NOD (n=5) and male NOR (n=4) mice with intermediate disease versus healthy male age-matched BALB/c (n=6) mice. Moderated t-statistics estimated using Limma R package. (.padj < 0.1, * padj < 0.05, ** padj < 0.01, *** padj < 0.001). *PL-7 – threonyl-tRNA synthetase; La; KU (P70/P80) – Heterodimer Ku protein subunits 70 and 80; GAD65 – Glutamic Acid Decarboxylase 65; LKM1 – Liver kidney microsome type 1;*

**Supplemental Figure 5.**
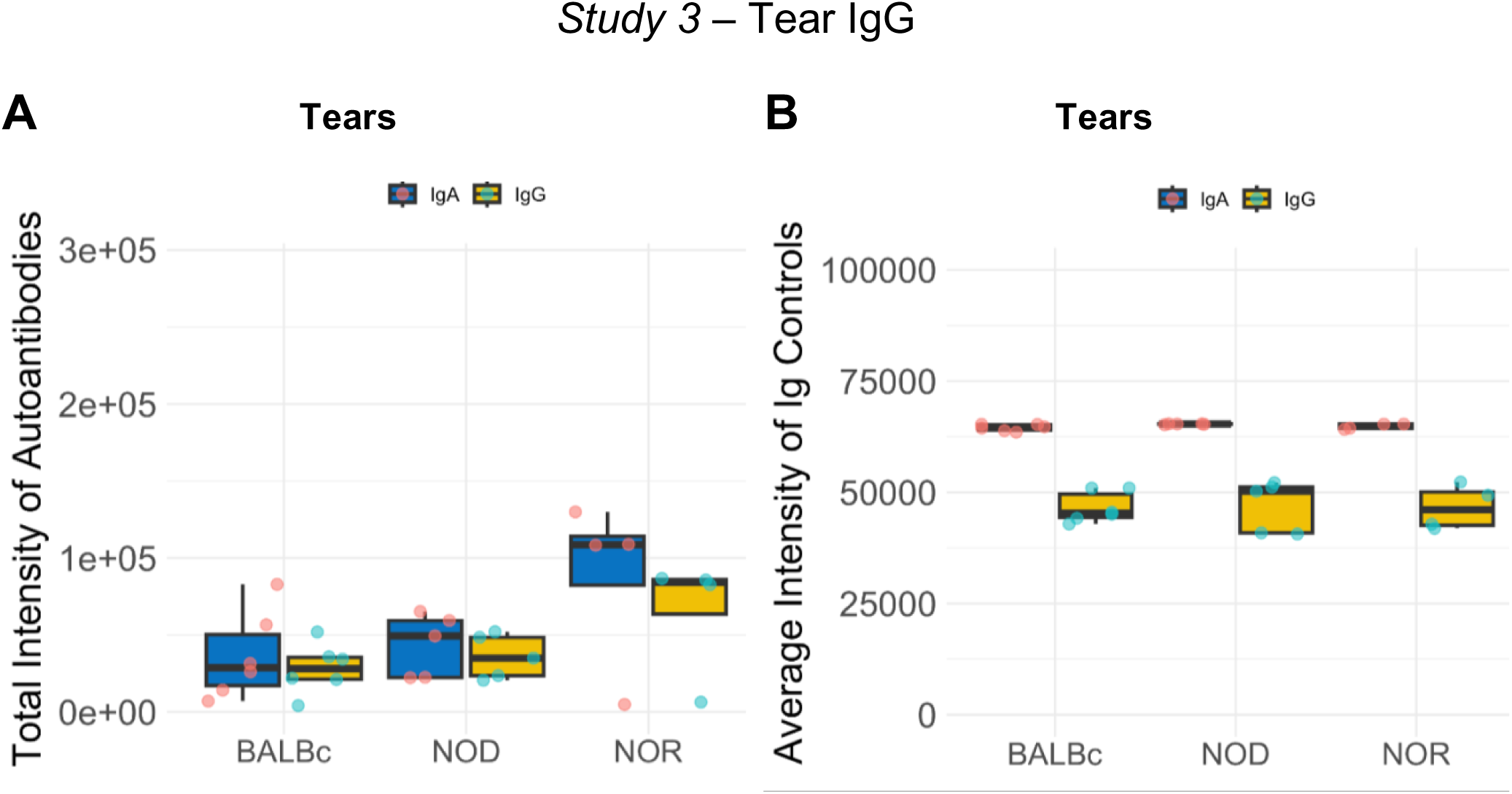
IgG vs IgA signal intensity in mouse tears from *Study 3*. **(A)** Boxplots showing sum of raw signal intensity of 80 autoantibodies for IgG and IgA in mouse tears. **(B)** Boxplots showing signal intensity of Ig reactivity towards mouse IgG or IgA control. The NOR and BALB/c mice with nearly 0 total signal intensity for both IgG and IgA in tears were removed from analysis. Each point represents one mouse, with serum and tears collected from male NOD (n=5) and male NOR (n=4) mice with intermediate disease versus healthy male BALB/c (n=6) mice.

**Supplemental Figure 6.**
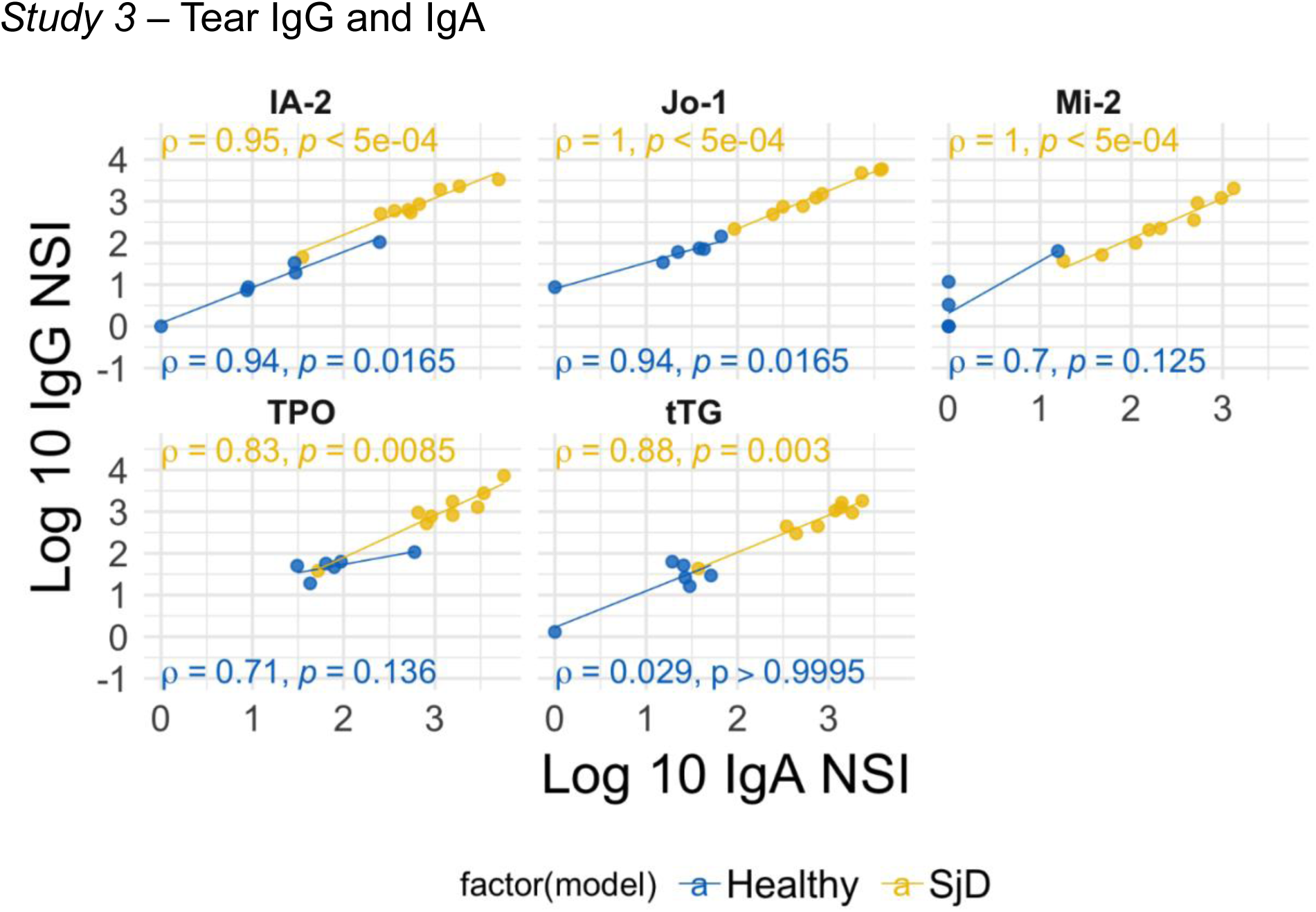
Correlation analysis of tear IgA vs IgG to the same autoantigen. Scatterplots showing log transformed raw intensity values of autoantibodies to the same autoantigen with significantly high reactivity in both IgA and IgG in both male NOD and NOR mice with intermediate disease with respect to male BALB/c. Each point represents one mouse. The male NOD and NOR mice are shown in yellow and healthy control BALB/c is shown in blue. Spearman’s correlation coefficients for tears from SjD mice (yellow) and healthy mice (blue) with respective p-values are also shown. Only p < 0.01 is considered significant to account to correct for multiple correlation. No mice were excluded from analysis.

**Supplemental Table 1.** Differentially Expressed IgG Autoantibodies in Serum *of Study 1* (NOD vs BALB/c).

|  | NOD (N=3) vs BALB/c (N=3)<br>Early Disease |  | NOD (N=3) vs BALB/c (N=3)<br>Advanced Disease |  | Age |
| --- | --- | --- | --- | --- | --- |
| Auto Antigen | P<sub>adj</sub> | Log Fold-Change<br>(Fold Change) | P<sub>adj</sub> | Log Fold-Change<br>(Fold Change) | P<sub>adj</sub> |
| PL-7<sup>†</sup> | **1.93E-02** | 6.91 (1002.49) | 0.10 | 3.82 (45.50) | 0.94 |
| PM/ScI-100 | **4.03E-04** | 6.29 (540.77) | **1.54E-03** | 4.92 (136.63) | 0.12 |
| La/SSB<sup>†</sup> | **3.45E-02** | 4.56 (95.40) | 0.083 | 3.31 (27.33) | 0.56 |
| LC1 | **1.78E-03** | 4.45 (85.66) | **1.54E-03** | 4.61 (100.11) | 0.56 |
| MAG-Fc | 0.065 | 3.57 (35.60) | **1.56E-02** | 6.19 (487.92) | 0.68 |
| KU (P70/P80) | **1.93E-02** | 2.88 (17.78) | **1.79E-02** | 2.87 (17.63) | 0.87 |
| PCNA | **5.05E-02** | 2.47 (11.82) | **1.03E-02** | 4.17 (64.68) | 0.77 |
| Mitochondrial antigen | **3.51E-02** | 2.08 (8.00) | 0.19 | 1.10 (3.02) | 0.75 |
| TPO | **5.28E-03** | 1.49 (4.44) | 0.10 | 1.06 (2.89) | - |
| Intrinsic Factor | **0.035** | 4.31 (74.44) | 0.40 | 1.37 (3.93) | - |
| CENP-B | 6.52E-02 | 1.58 (4.87) | - | - | **0.02** |
| SRP54<sup>></sup> | 5.05E-02 | -0.58 (1.78) | 0.03 | -0.70 (2.01) | **0.12** |

Mice with early disease were from 8-12 weeks while mice with advanced disease were >20 weeks as in Methods. BALB/c mice were age-matched to NOD mice
$p_{adj} < 0.05$ was considered significant to account for multiple comparisons. Statistically significant $p$ -values with fold change greater than 2 are shown in bold.
<sup>†</sup>Also upregulated in tears of male NOD and NOR mice compared to healthy BALB/c
<sup>></sup>IgG downregulated in tears of male NOD and NOR mice compared to healthy BALB/c

**Supplemental Table 2.** Differentially Expressed IgG Autoantibodies in Serum of *Study 2* (NOR vs BALB/c).

| Auto Antigen | P Value | Log<sub>2</sub> Fold-Change<br>(Fold Change) |
| --- | --- | --- |
| LC1 | **0.0012** | 4.24 (18.89) |
| PM/Scl 100 | **3.05 x 10<sup>-5</sup>** | 6.59 (96.33) |
| RPP P0 | **0.0015** | 4.13 (17.50) |
| Mito antigen | **0.0099** | 3.08 (8.45) |
| La/SSB<sup>†</sup> | **3.10 x 10<sup>-5</sup>** | 6.58 (95.67) |
| C3 | **0.0021** | 3.94 (15.35) |
| C4 | **0.0004** | 4.95 (30.91) |
| C5 | **0.0017** | 4.05 (16.56) |
| CRP | **0.0053** | 3.42 (10.70) |
| >M2 | **0.0001** | 5.67 (50.91) |
| Sm RNP | **0.0002** | 5.44 (43.41) |
| >SRP54 | **0.0016** | 4.08 (16.91) |
| TNF-α | **0.0053** | 3.42 (10.70) |
| Intrinsic Factor | **0.0159** | 2.82 (7.06) |
| PL-7<sup>†</sup> | **0.0201** | 2.69 (6.45) |
| PCNA | **0.0341** | 2.40 (5.27) |

*p* < 0.05 was considered significant to account for multiple comparisons.
Statistically significant *p*-values are shown in bold.
<sup>†</sup>Also upregulated in tears of male NOD and NOR mice compared to healthy BALB/c > IgG downregulated in tears of male NOD and NOR mice compared to healthy BALB/c

**Supplemental Table 3.** Differentially Expressed IgA Autoantibodies in Serum from *Study 3* (NOD, NOR vs BALB/c).

|  | NOD v BALB/c |  | NOR v BALB/c |  |
| --- | --- | --- | --- | --- |
|  | padj | Log Fold Change | padj | Log Fold Change |
| IF | **1.15e-03** | 4.77 (117.91) | **2.36e-06** | 8.10 (3.294x10<sup>3</sup>) |
| ssDNA | **0.0119** | 3.67 (39.25) | **7.64e-04** | 5.24 (188.67) |
| CytC | **3.98e-03** | 4.5 (90.01) | 0.1151 | 2.01 (7.46) |
| GP2 | 0.101 | 1.74 (5.70) | **5.59e-03** | 3.23 (25.27) |
| Gliadin | **3.49e-03** | 2.77 (15.96) | 0.0903 | 1.79 (5.99) |

