## Supplemental Methods 1 for "Serum and Tear Autoantibodies from NOD and NOR Mice as Potential Diagnostic Indicators of Local and Systemic Inflammation in Sjögren’s Disease"

#### Study 3 - Tear IgA Analysis

Shruti Kakan

Last Updated 2024-10-24

##### Loading Libraries

```
knitr::opts_chunk$set(
    echo = TRUE,
    message = FALSE,
    warning = FALSE,
    root.dir = "~/Documents/3_Parkinsons_disease/Autoantibody_Data/Tear_Auto_Validation_20
22/"
)
library(car)
library(lsmeans)
library(calibrate)
library(dplyr)
#library(DEGreport)
library(DESeq2)
library(DEFormats)
library(edgeR)
library(ggpubr)
library(ggsci)
library(ggplot2)
library(gridExtra)
library(pheatmap)
library(reshape2)
library(RColorBrewer)
library(scales)
library(rstatix)
library(tidyr)
library(magrittr)
#library(PCAtools)
library(tidyverse)
library(Biobase)
#library(marray)
library(limma)
library(gplots)

library(devtools)
#install_github("dpgaile/AutoAntArrayExmpl")
library(AutoAntArrayExmpl)

library(devtools)
#install_github("dpgaile/AutoAntArrayExmpl")
#devtools::install_github('renozao/NMF@devel')
library(AutoAntArrayExmpl)
library(NMF)
library(quantreg)
library(asbio)
library(fdrtool)
#library(discreteMTP)
library(scales)
library(ggsci)
library(ggplot2)
```

```
knitr::opts_chunk$set(
    echo = TRUE,
    message = FALSE,
    warning = FALSE,
    root.dir = '~/Documents/3_Parkinsons_disease/Autoantibody_Data/Tear_Auto_Validation_20
22/'
)

setwd('~/Documents/3_Parkinsons_disease/Autoantibody_Data/Tear_Auto_Validation_2022/')
#IgA_NSI <- read.csv("IgA_data/IgA_MCF_SSK_546_Tear_NSI_norm.csv",  header=T)[1:80,]
IgA_NSI <- read.csv("IgA_data/IgA_MCF_SSK_546_Tear_NSI.csv",  header=T)[1:80,]
IgA_SNR <- read.csv("IgA_data/IgA_MCF_SSK_546_Tear_SNR.csv",  header=T)[1:80,]
Strain <- c(rep("NOD", each=5),rep("NOR", each=3), rep("BALBc", each=6))
```

### Adding Auto-antigen id names

```
#Adding Auto-antigen id names
Antigen_ID <- read.csv("~/Documents/3_Parkinsons_disease/Autoantibody_Data/Tear_Auto_Valid
ation_2022/Antigen_ID.csv",  header=T)[1:80,1:2]

#colnames(IgA_NSI)[1] <- colnames(Antigen_ID)[2]
#colnames(IgA_SNR)[1] <- colnames(Antigen_ID)[2]

Antigen_ID[68,"ID"] <- IgA_NSI[68,"ID"]
IgA_NSI <- full_join(IgA_NSI, Antigen_ID, by="ID")
IgA_SNR <- full_join(IgA_SNR, Antigen_ID, by="ID")
rownames(IgA_NSI) <- IgA_NSI$Antigen_ID
rownames(IgA_SNR) <- IgA_SNR$Antigen_ID

IgA_NSI <- IgA_NSI[,-c(1, 17)]
IgA_SNR <- IgA_SNR[,-c(1, 17)]
```

#### Setting up column Metadata

```
Strain <- c(rep("NOD", each=5),rep("NOR", each=4), rep("BALBc", each=6))

colData <- as.data.frame(cbind(c(colnames(IgA_NSI)), Strain))
colnames(colData) <- c('Sample', "Strain")
rownames(colData) <- colData$Sample
colData$Strain <- factor(colData$Strain)
#colData$Strain <- relevel(colData$Strain, ref = "BALBc")
#Biofluid <- c( rep("Tear", each=11), rep("Serum", each=11))

#colData <- as.data.frame(cbind(c(colnames(IgA_NSI)), Strain, Biofluid))
```

### Filtering Data based on low Signal to Noise ratio

Rows with low signal to noise ratio of less than rowmeans 2.8 were removed. Additionally rows with low overall signal intensity were also removed.

```
IgA_raw=list()
IgA_SNR$average <- rowMeans(as.matrix(IgA_SNR))
IgA_SNR$med <- rowMedians(as.matrix(IgA_SNR))

IgA_raw$NSI <- as.matrix(IgA_NSI[which(IgA_SNR$average>2.8),])
IgA_raw$SNR <- as.matrix(IgA_SNR[which(IgA_SNR$average>2.8),])[,1:15]
#IgA_raw$NSI <- as.matrix(IgA_NSI[which(rowSums(IgA_SNR[,1:15]>2.8) > 6),])
#IgA_raw$SNR <- as.matrix(IgA_SNR[which(rowSums(IgA_SNR[,1:15]>2.8) > 6),][,1:15])

log2(colSums(IgA_raw$NSI+0.5))
```

```
##     NOD_M1_Tears     NOD_M3_Tears     NOD_M4_Tears      NOD_M5_Tear
##         15.58217         15.99049         14.43786         15.85415
##      NOD_M6_Tear     NOR_M1_Tears NOR_M2_SSK_Tears     NOR_M3_Tears
##         14.43494         16.98300         16.72778         12.24237
##     NOR_M4_Tears   BALBc_M1_Tears   BALBc_M2_Tears   BALBc_M3_Tears
##         16.70994         14.66195         12.77064         15.78438
##   BALBc_M4_Tears   BALBc_M5_Tears   BALBc_M6_Tears
##         16.33585         14.93757         13.78248
```

This reduced the total number of Autoantibodies (rows) included in the analysis from 80 to 69.

### Visualizing Filtered Data

```
dataN <- log2(IgA_NSI + 0.5)
countData = as.data.frame(dataN)

df_dseq = melt(countData, variable.name = "Samples", value.name = "count")# reshape the ma
trix

mycolors <- colorRampPalette(brewer.pal(8, "Set1"))(15)

ggplot(df_dseq, aes(x = count, color=Samples)) +
   geom_density(alpha = 0.5, size = 0.8) +
   #facet_wrap(~Strain, ncol=2) +
   theme_minimal() + #xlim(-1.5,6) +
   scale_colour_manual(values=mycolors, name="") +
   guides(fill="none")
```

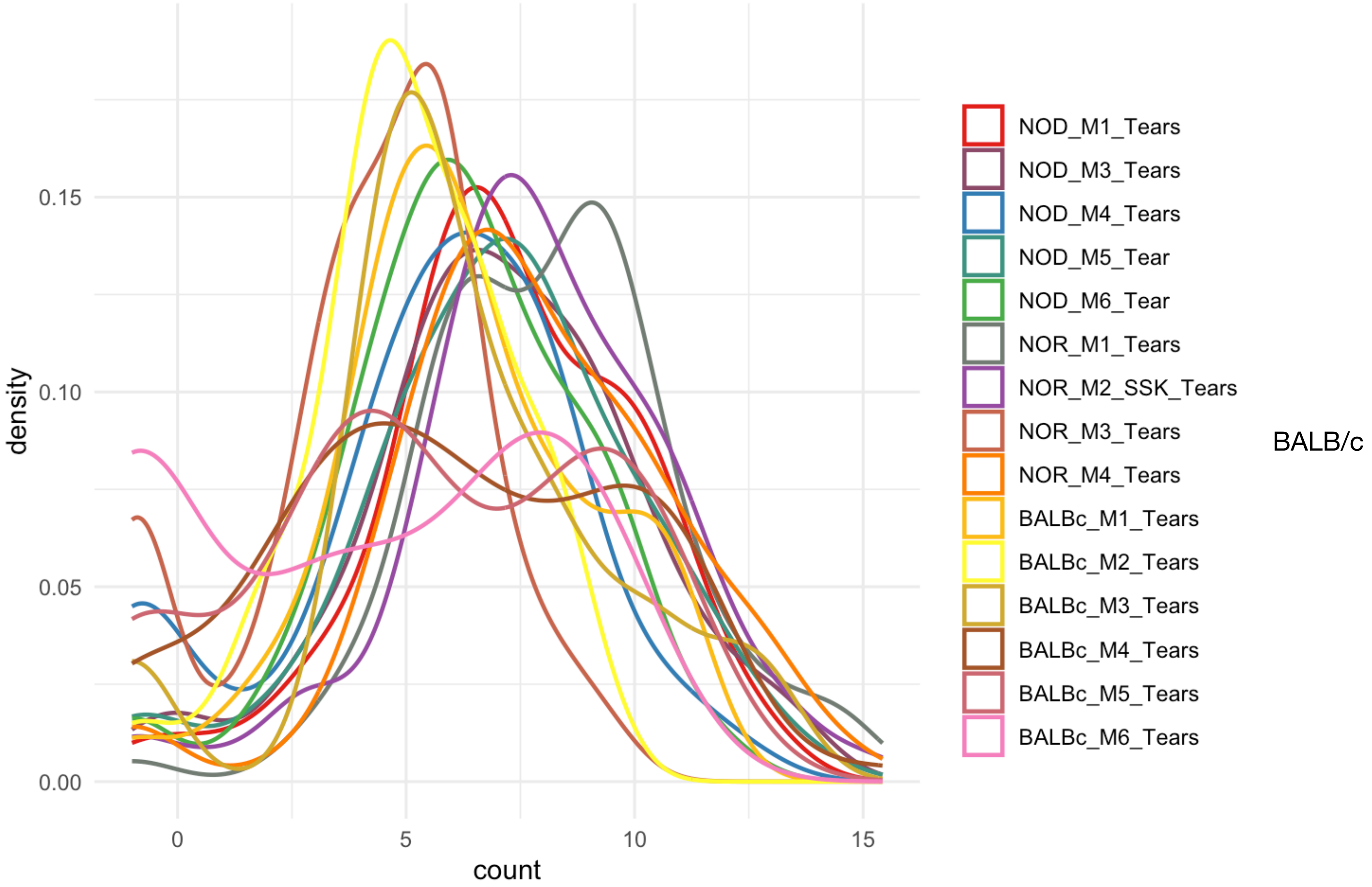


Sample M6 appears to have extremely low signal intensity values across the board.

```
boxplot(as.data.frame((dataN)),main="Tear IgA Signal Intensity Prior to Row filtering")#,col=Sample)
```

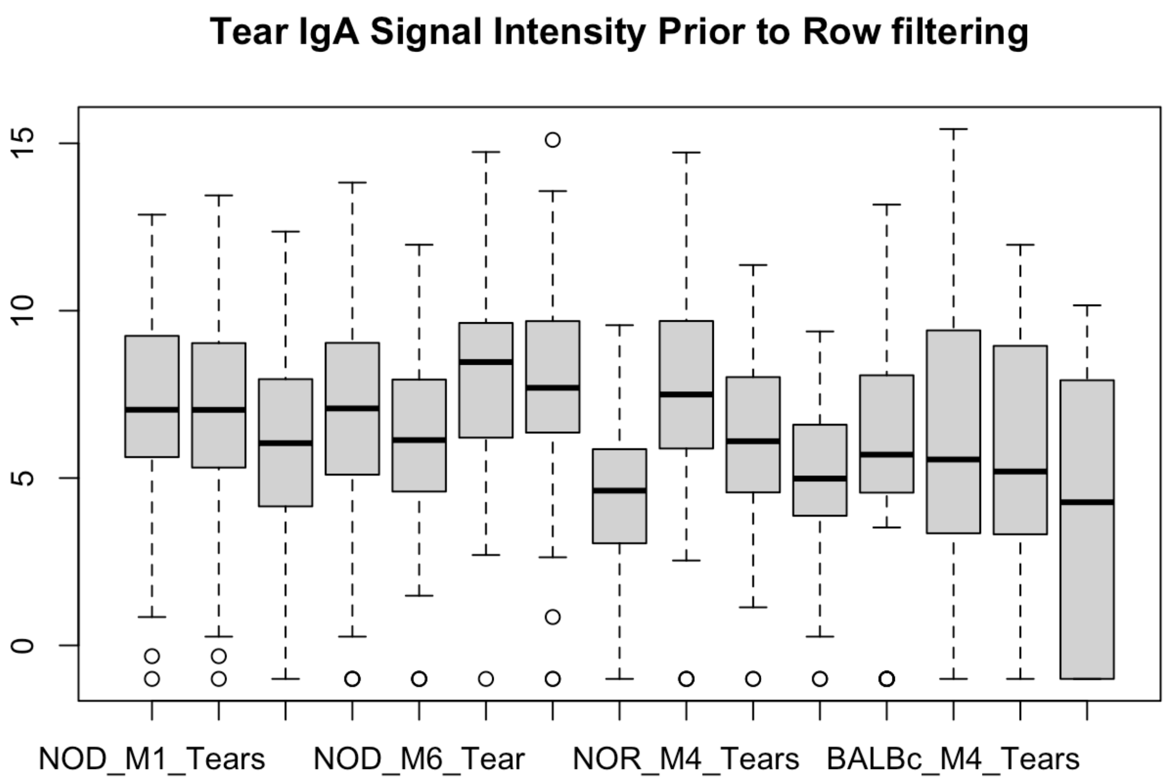


```
#dataN <- (IgA_NSI * IgA_SNR[,1:15] )
dataN <- (IgA_NSI)
countData = as.data.frame(dataN)
#boxplot(as.data.frame((dataN)),main="NSI")

#dataN <- log2(IgA_raw$NSI * IgA_raw$SNR +0.5)
dataN <- log2(IgA_raw$NSI+0.5)
countData = as.data.frame(dataN)
boxplot(as.data.frame((dataN)),main="Tear IgA Signal Intensity After Row filtering")
```

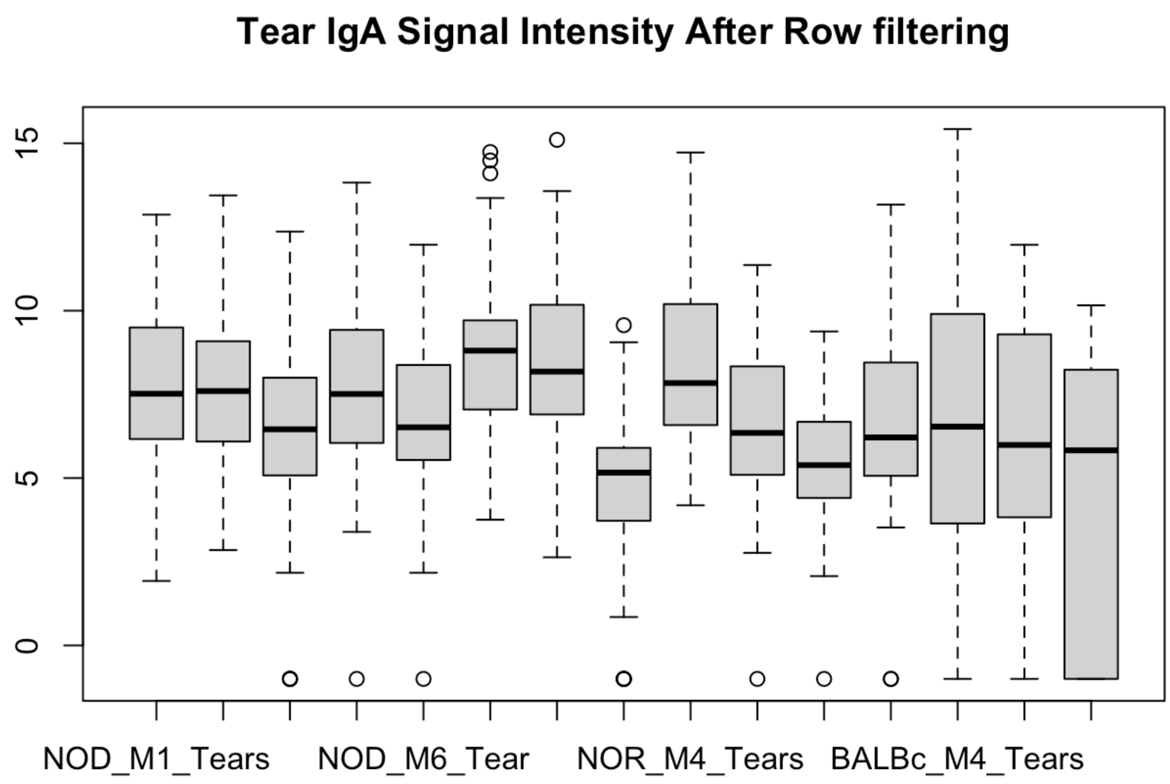


### Limma Based Default Normalization & DE Analysis

Sample BALB/c M6 was excluded from normalization and differential expression analysis.

```
library(limma)
#dataN <- log2(IgA_raw$NSI * IgA_raw$SNR +0.5)[,-15]
dataN <- log2(IgA_raw$NSI +0.5)[,-15] #Sample BALB/c M6 removed
mydata <- as.matrix(dataN)

conditions<- paste(colData$Strain[-15],sep=".")
conditions <- factor(conditions, levels=unique(conditions))
design <- model.matrix(~0+ conditions)
colnames(design) <- levels(conditions)
fit <- lmFit(mydata, design)

cont.matrix<- makeContrasts(
   NTvBT =  NOD - BALBc,
   nTvBT = NOR - BALBc,
   levels = design)
fit.cont<- contrasts.fit(fit, cont.matrix)
fit.cont<- eBayes(fit.cont)

#MA plots showing relationship between average expression of autoantibodies and their log
fold-changes
plot.new()
plotMD(fit.cont, col=1) # NOD v BALB/c
```

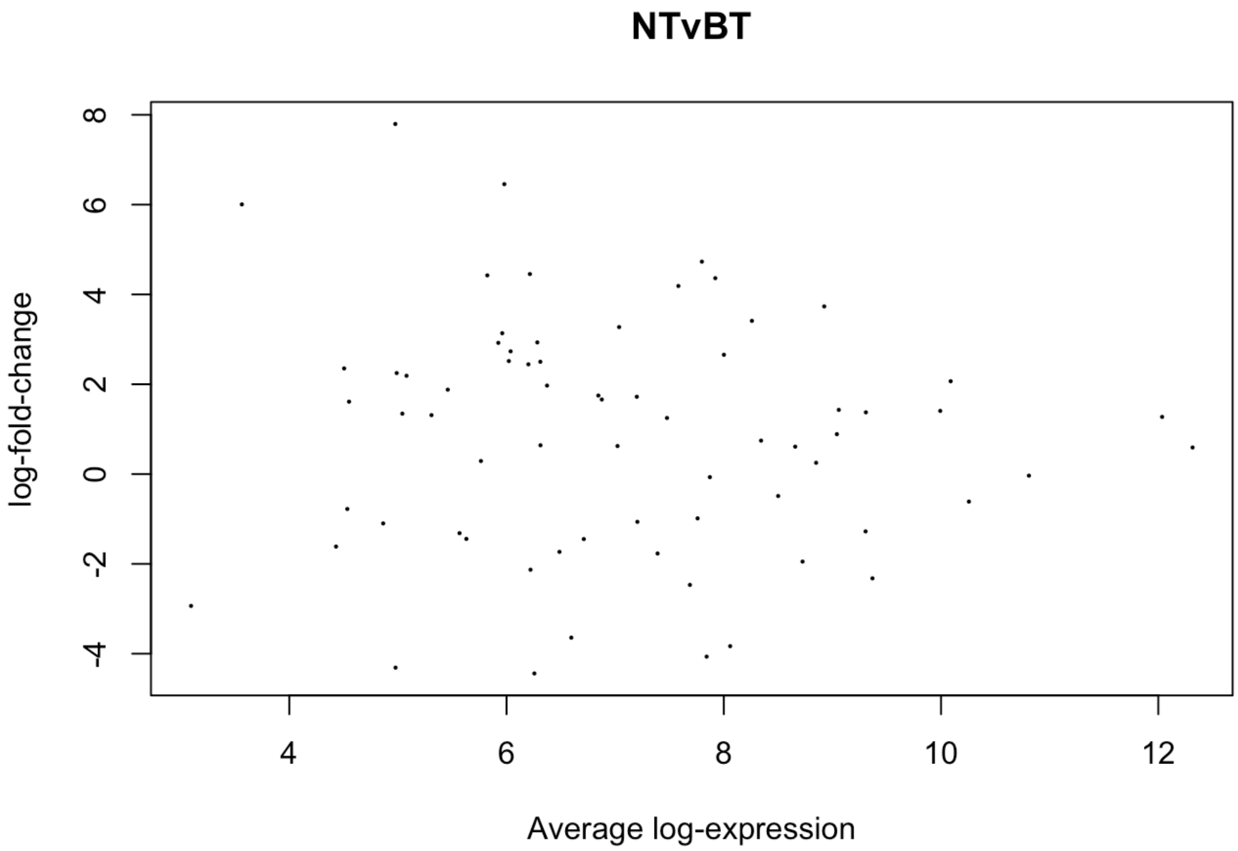


```
plotMD(fit.cont, col=2) # NOR v BALB/c
```

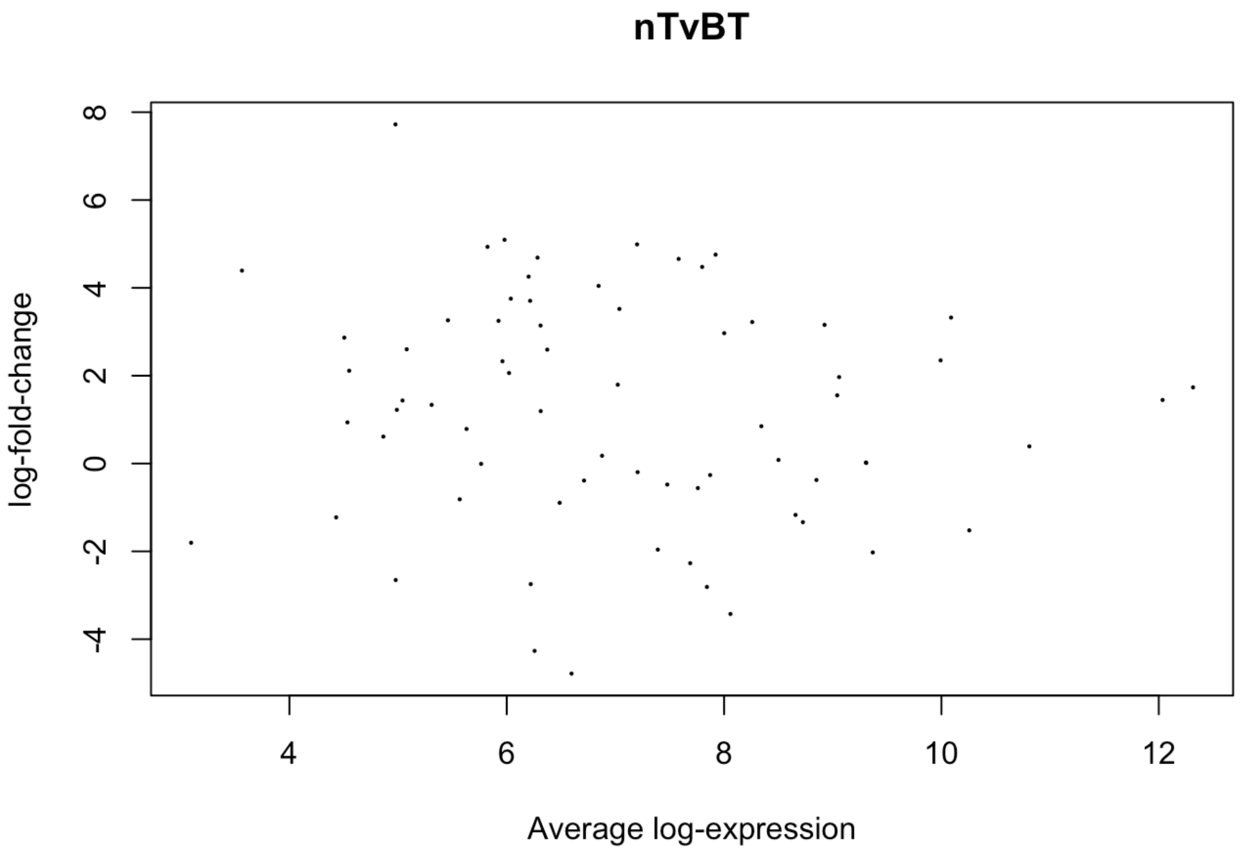


```
# QQ Plots of normalized data
qqt(fit.cont$t,df=fit.cont$df.prior+fit.cont$df.residual)
```

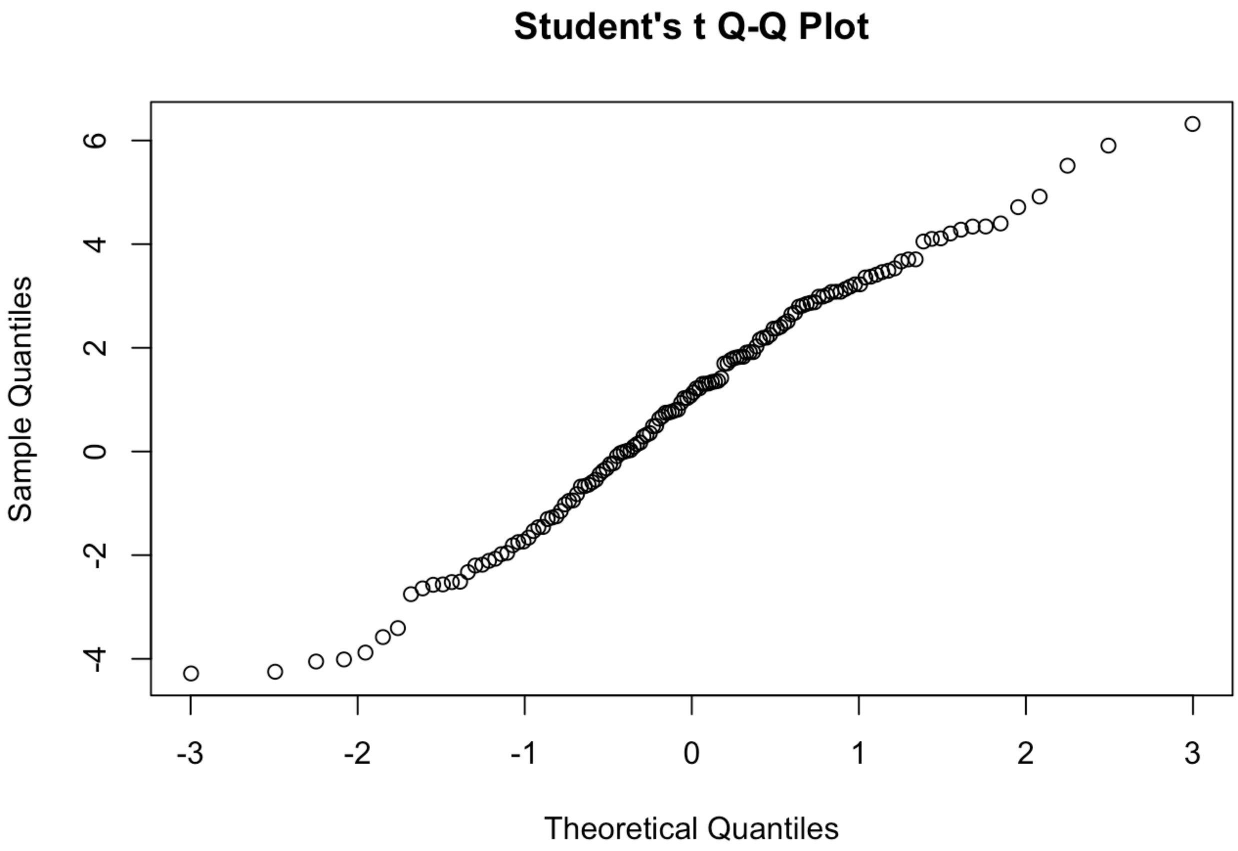


```
#Density plots of normalized Data
plotDensities(fit)
```

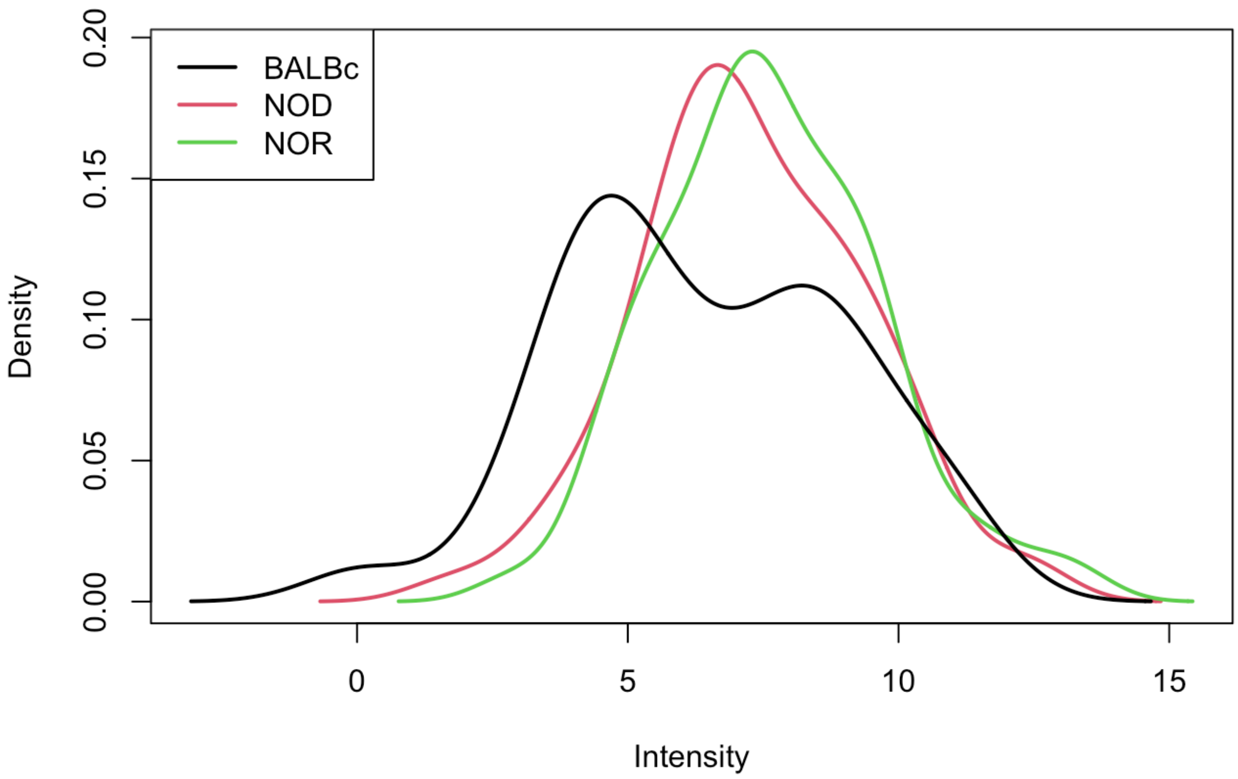


While this normalization works fairly well, most of the BALB/c samples show a relatively low signal intensity across the board. While this my be a true biological effect, we have to consider it as a potential batch effect owing to the fact that BALB/c is a closely matched control of NOD and NOR.

```
#Generating results data frames using Limma's topTable() function
setwd("~/Documents/3_Parkinsons_disease/Autoantibody_Data/Tear_Auto_Validation_2022/IgA_da
ta/")
NTvsBT <- topTable(fit.cont, coef=1, p.value=1, number=50, adjust.method = 'BH')
nTvsBT <- topTable(fit.cont, coef=2, p.value=1, number=50, adjust.method = 'BH')
NTvsBT$Antigen <- row.names(NTvsBT)
nTvsBT$Antigen <- row.names(nTvsBT)
NTvnTvBT <- full_join(NTvsBT, nTvsBT,
                      by="Antigen",
                      suffix = c(".NOD", ".NOR"))
rownames(NTvnTvBT) <- NTvnTvBT$Antigen
```

### Results Table

```
#Generating output file
NTvnTvBT[is.na(NTvnTvBT)] <- 0.5
NTvnTvBT <- NTvnTvBT[-c(which(NTvnTvBT$adj.P.Val.NOD > 0.1 & NTvnTvBT$adj.P.Val.NOR > 0.
1)),]
write.csv(NTvnTvBT, file="Tear_IgA_Limma.csv", sep=',')

print(NTvnTvBT[which(NTvnTvBT$logFC.NOD>1),c(1:2,5,8,12)])
```

```
##             logFC.NOD AveExpr.NOD adj.P.Val.NOD logFC.NOR adj.P.Val.NOR
## Mi-2         7.795830    4.975896  0.0002787797  7.722650   0.000683158
## SAE1 SAE2    6.455780    5.979389  0.0008008029  5.094235   0.005698873
## tTG          4.730122    7.798166  0.0032457358  4.476692   0.005224389
## Jo-1         4.362181    7.921660  0.0047276686  4.756630   0.004607082
## LKM 1        4.453778    6.214859  0.0049074651  3.705070   0.017644014
## TNF a        6.004939    3.563230  0.0049074651  4.392796   0.032703291
## IA-2         4.187879    7.581873  0.0134489150  4.659095   0.009911925
## KS           4.424495    5.822803  0.0134489150  4.934300   0.009911925
## GAD65        3.411583    8.258565  0.0134489150  3.221565   0.020878584
## TPO          3.732875    8.924370  0.0158256914  3.157495   0.040459601
## Gliadin      3.272999    7.036221  0.0221382634  3.520608   0.017644014
## PM Scl75     3.136187    5.960308  0.0259014278  2.328724   0.091275690
## KU P70P80    2.921768    5.924015  0.0299842041  3.248915   0.020472567
## La SSB       2.932182    6.283174  0.0348995668  4.688158   0.004607082
## PL-7         2.656189    8.000905  0.0348995668  2.967767   0.022646584
## LC1          2.500449    6.310963  0.0348995668  3.141955   0.014259124
## IF           2.443567    6.200971  0.0494558660  4.255189   0.004607082
## PCNA         2.188723    5.079314  0.0555662482  2.601838   0.032513431
## SmD1         2.516116    6.020889  0.0582921581  2.061221   0.132603261
## PL-12        1.970215    6.372643  0.0644397656  2.593297   0.023301083
## Cardiolipin  2.732643    6.037356  0.0662231843  3.753691   0.020878584
## GP2          2.067020   10.088505  0.0831644166  3.324054   0.014121733
## BPI          1.611897    4.550018  0.1379397699  2.112938   0.069630435
## NXP2         1.721346    7.199159  0.1588001089  4.989552   0.003059614
## GP210        1.748597    6.845041  0.1725706594  4.044152   0.009911925
## C3           1.879431    5.458823  0.2870342822  3.261196   0.089106039
```

16 IgA autoantibodies are significantly upregulated in both NOD & NOR as compared to male BALB/c with adjusted p value < 0.05

### Voom Normalization After Removing outliers

Using the Limma Voom package and a quantile normalization approach, we hope to completely remove the batch effect. This should give us a more conservative result.

```
#After removing outlier samples BALB/c 6

###### Voom normalization with quantiles
conditions<- paste(colData$Strain[-15],sep=".")
conditions <- factor(conditions, levels=unique(conditions))
design <- model.matrix(~0+ conditions)
colnames(design) <- levels(conditions)

v <- voom(counts=(IgA_raw$NSI[,-15]+0.5), design, plot=TRUE, normalize="quantile")
```

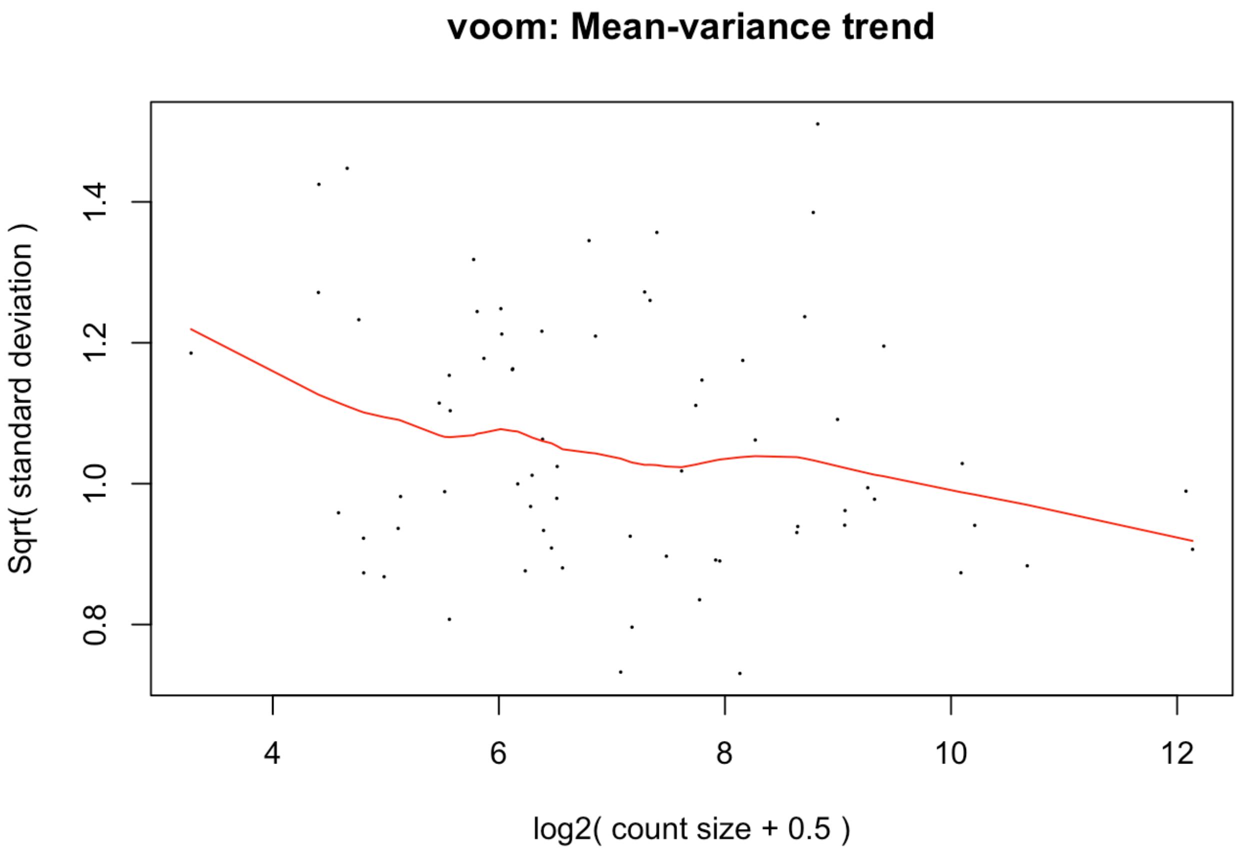


```
#v <- voom(dae, design, plot=TRUE)
fit <- lmFit(v, design)
fit.cont<- contrasts.fit(fit, cont.matrix)
fit.cont<- eBayes(fit.cont)
#plot(x=colSums(IgA_raw$NSI), y=colSums(IgG_raw$NSI))
```

#### QC plots of analysis

```
#PC1 separates Diseases and healthy samples
plotMDS(v,col=as.numeric(Strain))
```

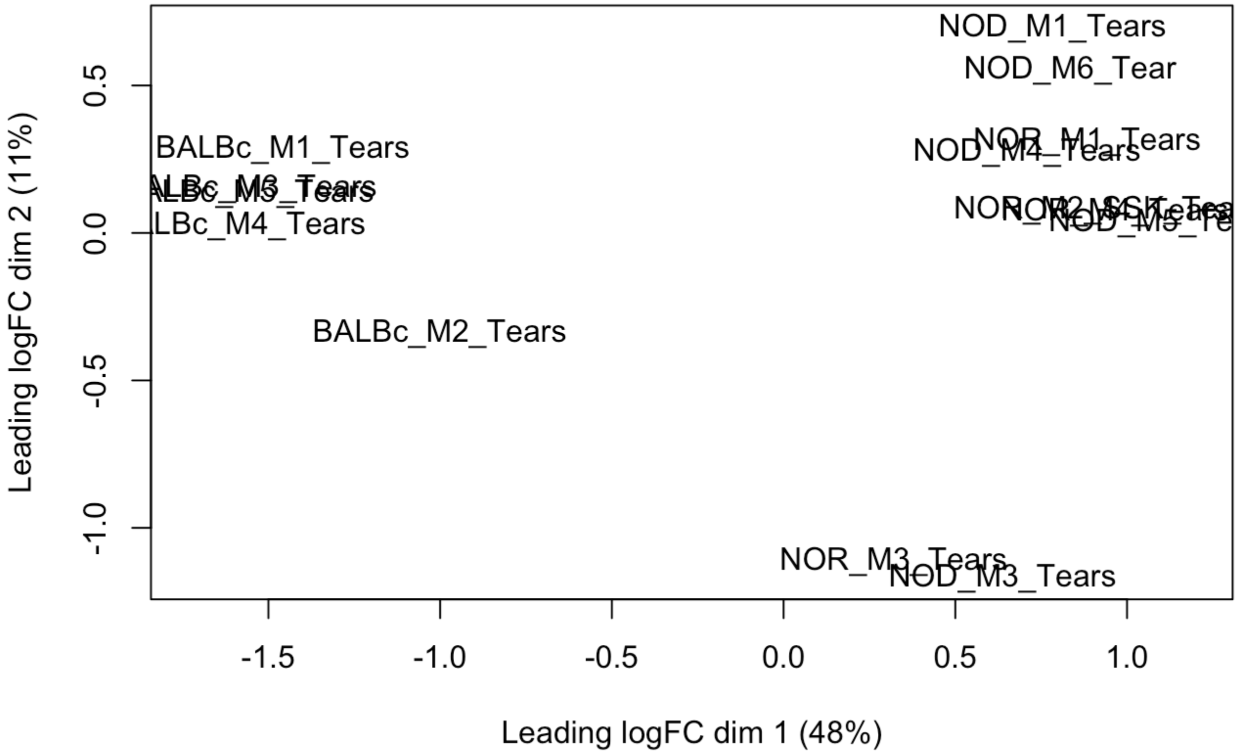


```
#MA plots showing relationship between average expression of autoantibodies and their log fold-changes
plot.new()
plotMD(fit.cont, col=1) # NOD v BALB/c
```

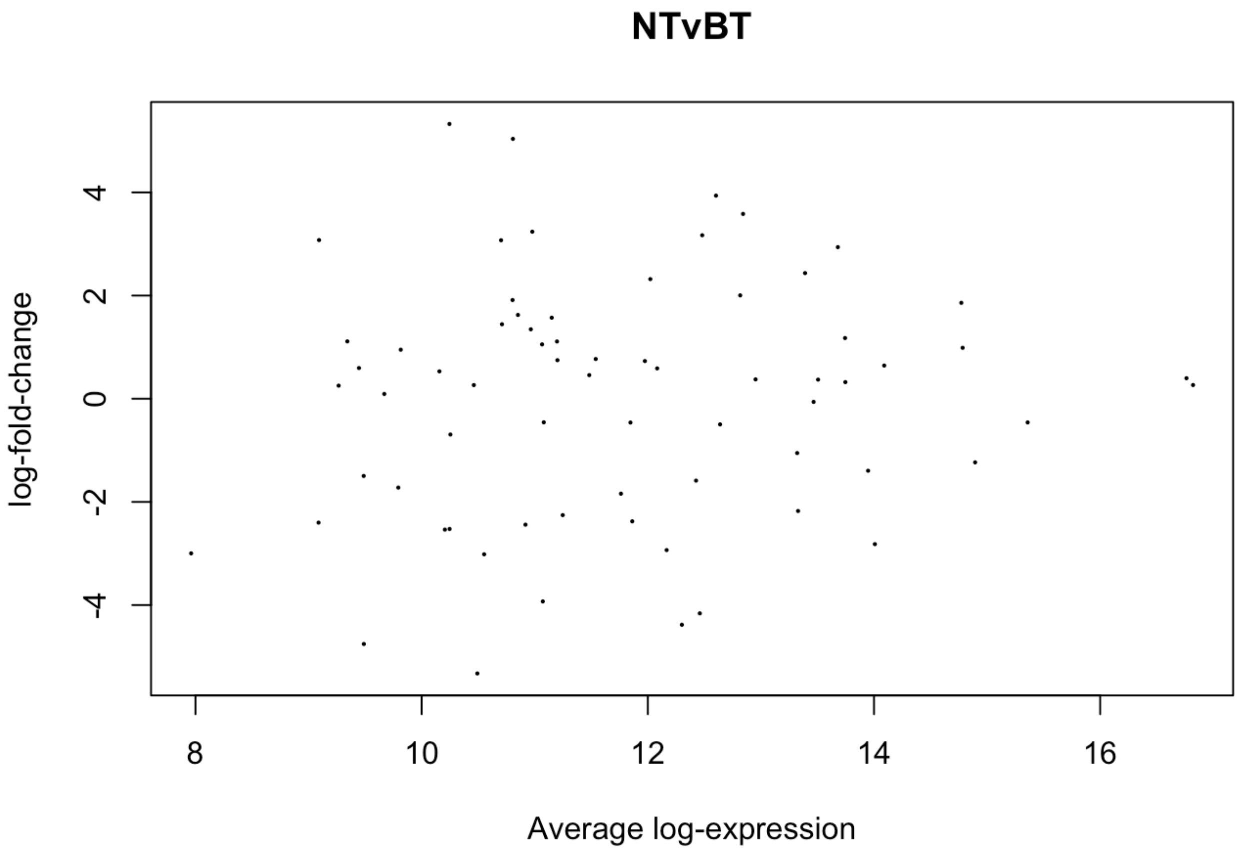


```
plotMD(fit.cont, col=2) # NOR v BALB/c
```

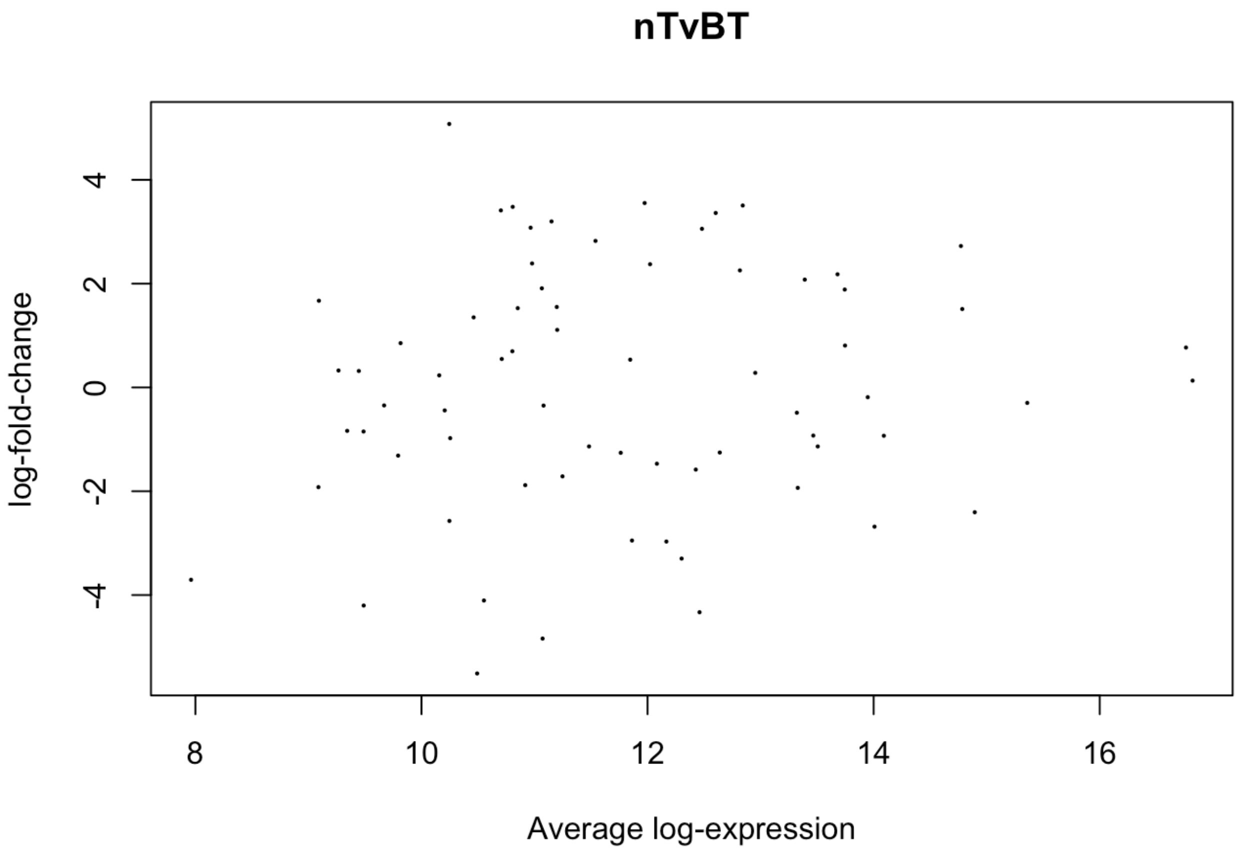


```
# The Q-Q plot is a fairly straight line
qqt(fit.cont$t,df=fit.cont$df.prior+fit.cont$df.residual)
```

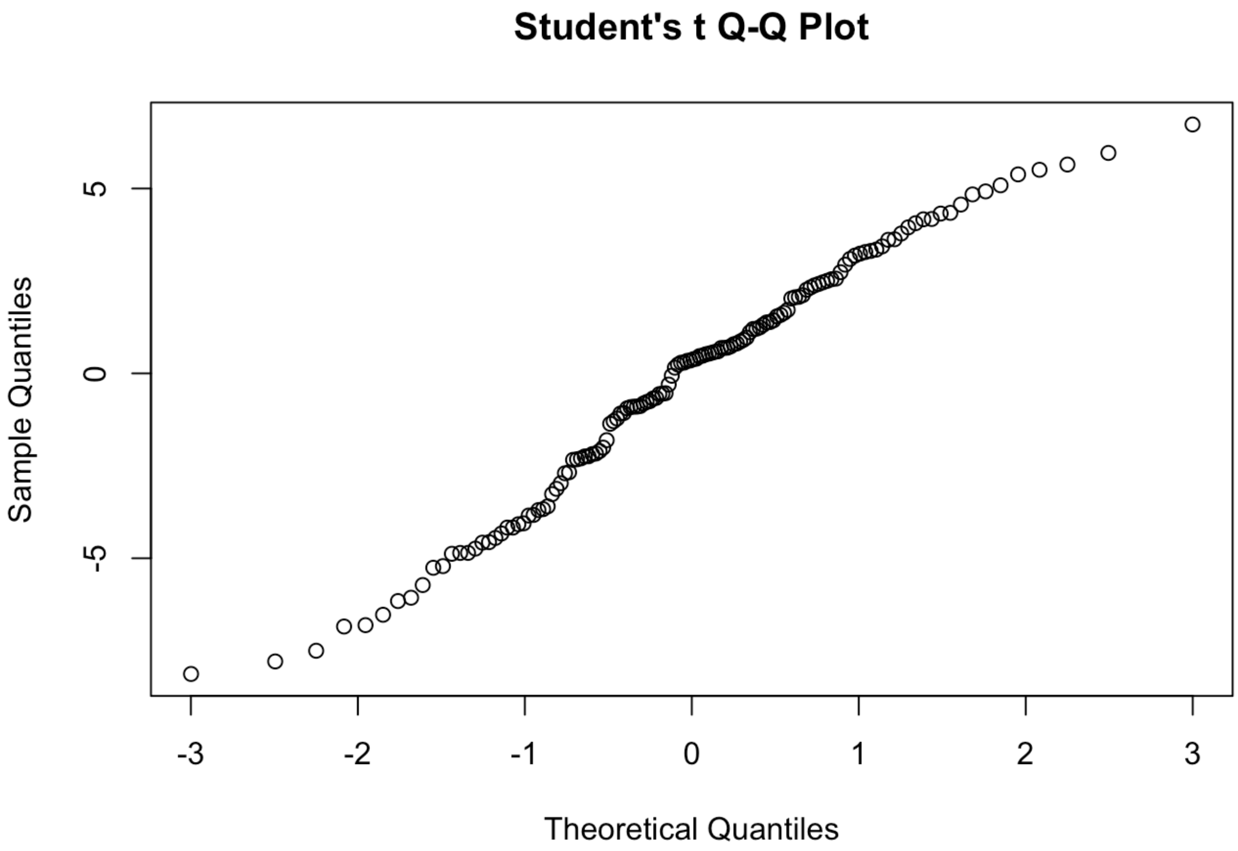


```
# The Density plots of all three groups overlap fairly well, confirming that the batch eff
ect has been largely normalized
plotDensities(v, group=Strain, col=c("orange","green", "blue"), log=TRUE)
```

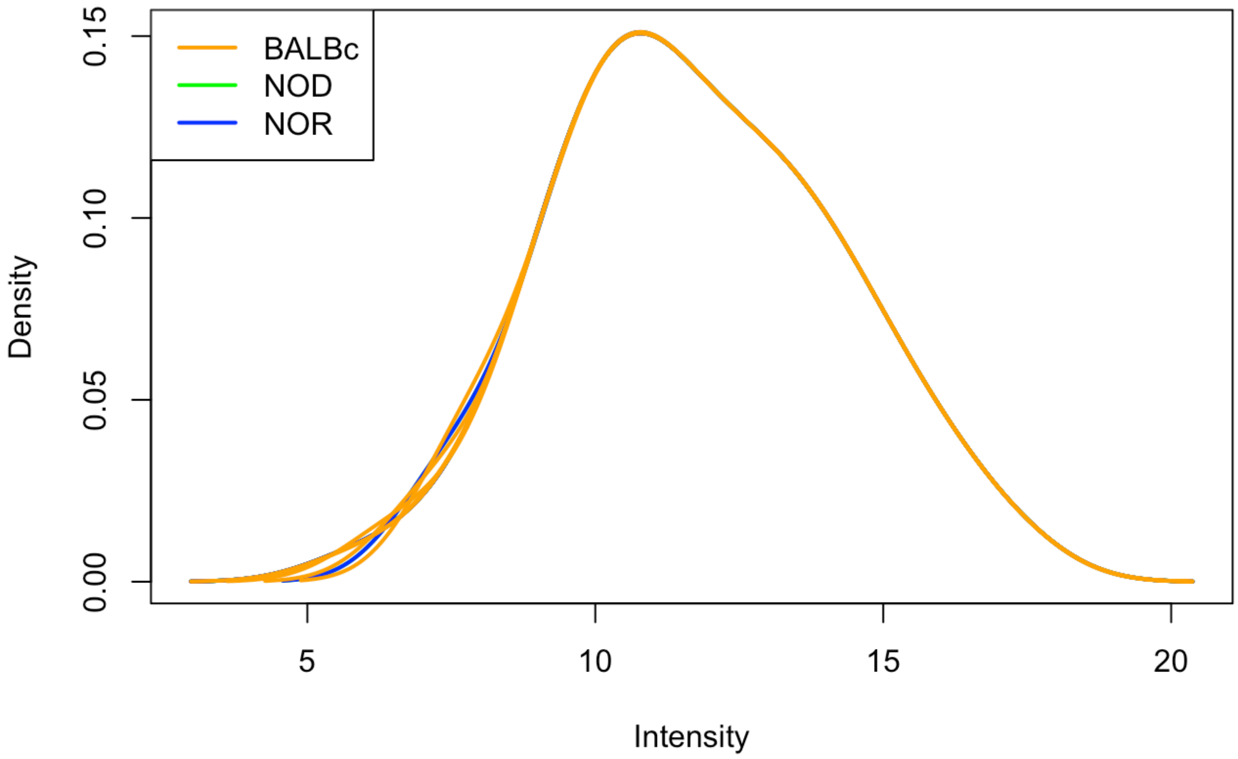


```
#Density plots

df_dseq = melt(v$E, variable.name = "Samples", value.name = "count")# reshape the matrix
df_dseq$Strain <- factor(substr(df_dseq$Var2, 1,3))
mycolors <- colorRampPalette(brewer.pal(8,"Set1"))(15)

ggplot(df_dseq, aes(x = count, color=Var2)) +
   geom_density(alpha = 0.5, size = 0.8) +
   facet_wrap(~Strain, ncol=2) +
   theme_minimal() + xlim(-5, 25) +
   scale_colour_manual(values=mycolors, name="") +
   guides(fill="none")
```

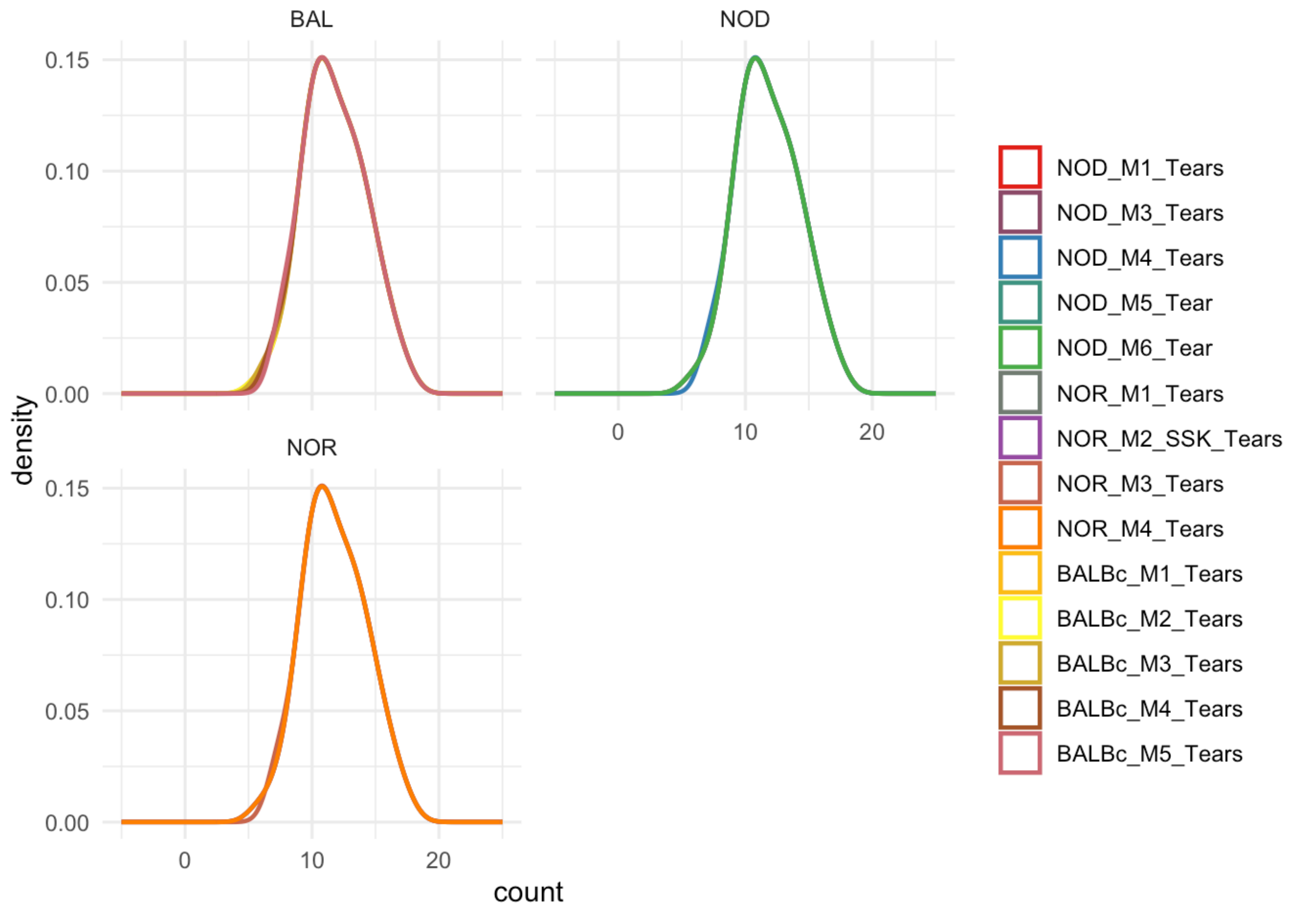


Sample wise Density plots facetted by the mouse strain show that voom normalized Log2 transformed NSI data of each sample overlap quite well and follow normal distribution.

### QC: Assessing Goodness of fit

```
#R-Squared ....... goodness of fit
for (i in 1:69){
   sst <- rowSums(v$E^2)
   ssr <- sst - fit.cont$df.residual*(fit.cont$sigma^2)
   Rsq<- (ssr/sst)
}
plot(1:69, Rsq)
```

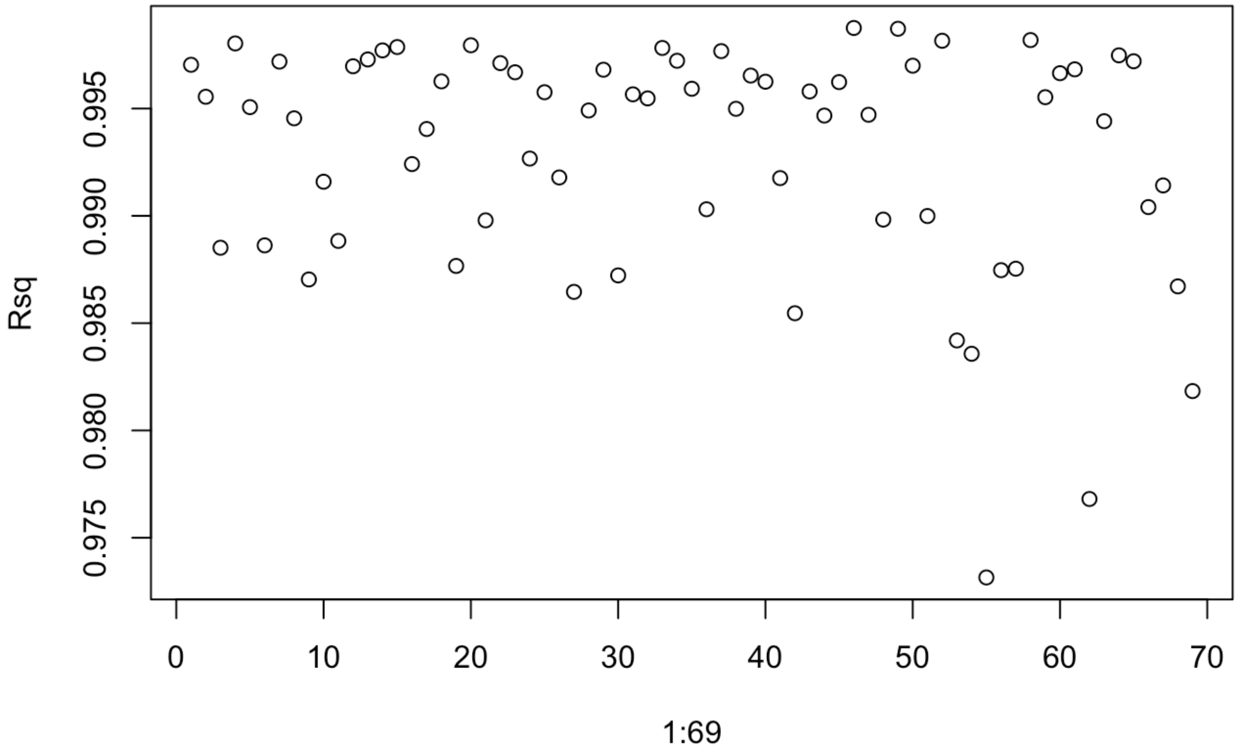


```
which(Rsq<0.90)
```

```
## named integer(0)
```

```
summary(fit.cont$r.squared)
```

```
## Length  Class   Mode
##      0   NULL   NULL
```

### Results Table: Limma-Voom (Quantile) Normalized DE

```
setwd("~/Documents/3_Parkinsons_disease/Autoantibody_Data/Tear_Auto_Validation_2022/IgA_da
ta/")
NTvsBT <- topTable(fit.cont, coef=1, p.value=1, number=50, adjust.method = 'BH')
nTvsBT <- topTable(fit.cont, coef=2, p.value=1, number=50, adjust.method = 'BH')
NTvsBT$Antigen <- row.names(NTvsBT)
nTvsBT$Antigen <- row.names(nTvsBT)
NTvnTvBT <- full_join(NTvsBT, nTvsBT,
                      by="Antigen",
                      suffix = c(".NOD", ".NOR"))
rownames(NTvnTvBT) <- NTvnTvBT$Antigen
NTvnTvBT[is.na(NTvnTvBT)] <- 0.5
NTvnTvBT <- NTvnTvBT[-c(which(NTvnTvBT$adj.P.Val.NOD > 0.1 & NTvnTvBT$adj.P.Val.NOR > 0.
1)),]
write.csv(NTvnTvBT, file="Tear_IgA_voom.csv", sep=',')
print(NTvnTvBT[which(NTvnTvBT$logFC.NOD>1),c(1:2,5,8,12)])
```

```
##             logFC.NOD AveExpr.NOD adj.P.Val.NOD logFC.NOR adj.P.Val.NOR
## tTG          3.937176   12.602754  3.011384e-05  3.361831  0.0003308796
## Mi-2         5.326484   10.245786  1.018836e-04  5.076768  0.0003639110
## SAE1 SAE2    5.037635   10.807100  1.513204e-04  3.479841  0.0056484908
## LKM 1        3.238098   10.978337  7.383427e-04  2.389129  0.0103068893
## Jo-1         3.582995   12.842062  1.652481e-03  3.505987  0.0019247068
## TPO          2.938684   13.680440  2.148782e-03  2.180326  0.0190203813
## IA-2         3.169350   12.481188  2.453302e-03  3.056493  0.0042021337
## PL-7         2.003696   12.816993  3.016223e-03  2.255281  0.0015144321
## GP2          1.859388   14.772166  7.465660e-03  2.725116  0.0005967557
## KS           3.072038   10.702153  1.265199e-02  3.411390  0.0098838751
## GAD65        2.435629   13.390596  1.495069e-02  2.077640  0.0434380159
## TNF a        3.075841    9.092553  3.188868e-02  1.670710  0.2882467679
## La SSB       1.571117   11.150277  4.190615e-02  3.198636  0.0005282556
## Gliadin      2.319357   12.022377  4.618957e-02  2.374340  0.0502645059
## KU P70P80    1.624575   10.851631  5.328293e-02  1.527862  0.0861559385
## PM Scl75     1.913631   10.803817  6.151310e-02  0.500000  0.5000000000
## IF           1.345973   10.965439  9.421284e-02  3.077987  0.0010620863
## dsDNA        1.176245   13.744256  9.795552e-02  1.886362  0.0108054878
## LC1          1.110343   11.196918  1.674744e-01  1.550423  0.0695410741
## Cardiolipin  1.053826   11.064888  3.625734e-01  1.909899  0.0897869926
```

Thirteen Auto-antibodies are significantly upregulated in tears of both male NOD & male NOR as compared to tears of healthy male BALB/c.

### Plotting DE Autoantibodies

#### DE Analysis using voom normalized counts

### Boxplots from voom normalized counts

```
chart_design <- theme(
   plot.title = element_text(color = "Black", size = 16, face = "bold", margin = margin(b=
15), hjust=0.4),
   axis.text.x = element_text(size=15),
   axis.text.y = element_text(size=14),
   axis.title.x = element_blank(),
   legend.text = element_text(size=15),
   legend.title = element_blank(),
   legend.position = "bottom",
   axis.title.y = element_text(size=18, margin = margin(r = 5)),
   strip.text.x = element_text(size =16, margin = margin(b=20), face='bold', hjust=0.4),
   strip.background = element_blank(),
   strip.placement = "outside")

mydata <- as.matrix(v$E)
hits <- rownames(NTvnTvBT)
hits[(length(hits)+1)] <- "Ro SSA 52"
#hits[(length(hits)+1)] <- "RoSSA 60"

Y=matrix(nrow=length(hits),ncol=14)

for (i in 1:length(hits)) {
   Y[i,] <- mydata[hits[i],]
}
rownames(Y) <- hits
colnames(Y) <- colData$Sample[-15]

Y <- as.data.frame(t(Y))
Y$Strain <- colData$Strain[-15]
Y$Sample <- paste0(Y$Strain, c(1:5,1:4, 1:5))

setwd("~/Documents/3_Parkinsons_disease/Autoantibody_Data/Tear_Auto_Validation_2022/IgA_da
ta/")
for (i in length(hits)){
   filename <- paste(hits[i],"IgA_Tears.tiff", sep="")
   p<- ggplot(Y, aes(x=Strain, y=Y[,i], fill=Strain)) +
      geom_boxplot(outlier.shape = NA, width = 0.5, coef=1, varwidth=F, show.legend = T, s
ize=0.9, position = position_dodge(0.9)) +
      geom_jitter(color = "darkgray", alpha =0.7, size=2.5, show.legend = F, position = po
sition_jitterdodge(dodge.width=0.9))+
      scale_color_manual(values=c("black", "navy")) +
      theme_minimal() +
      chart_design +
      ylab("Log Norm Intensity")  +
      scale_x_discrete(labels= c("BALB/c", "NOD", "NOR")) +
      labs(title=colnames(Y[i]), hjust=0.5) +
      scale_fill_jco()
   tiff(filename, units="in", width=3.45, height=3.5, res=300)
   print(p)
   dev.off()
```

```
}
#print(p)
```

#### Combined Boxplots of Upregulated hits

```
chart_design <- theme(
   #plot.title = element_text(color = "Black", size = 16, face = "bold", margin = margin(b
=15), hjust=0.4),,
   axis.text.x = element_text(size=16, margin = margin(b=5)),
   axis.text.y = element_text(size=18),
   axis.title.x = element_blank(),
   legend.text = element_blank(),
   legend.title = element_blank(),
   legend.position = "top",
   axis.title.y = element_text(size=24, margin = margin(r = 5)),
   strip.text.x = element_text(size =19, margin = margin(b=15), face='bold', hjust=0.4),
   strip.background = element_blank(),
   strip.placement = "outside")

#Generating Data Frame for plotting Combined boxplot
Y_combined <- Y[,c(NTvnTvBT$Antigen[which(NTvnTvBT$adj.P.Val.NOD<0.051 & NTvnTvBT$adj.P.Va
l.NOR < 0.051 & NTvnTvBT$logFC.NOD>0)])]
#Y_combined[,14:15] <- Y[,c("IF", "La SSB")]
Y_combined[,(ncol(Y_combined)+1):(ncol(Y_combined)+2)] <- Y[,(ncol(Y)-1):(ncol(Y))]
Y_combined <- gather(Y_combined, "Antigen", "V Counts", 1:(ncol(Y_combined)-2))
#Y_combined[which(Y_combined$Strain=="BALBc"),"Strain"] = factor("BALB/c")



#Generating combined boxplot of 13 autoantibodies using ggplot()
setwd("~/Documents/3_Parkinsons_disease/Autoantibody_Data/Tear_Auto_Validation_2022/IgA_da
ta/")
tiff("Tear_IgA_hits.tiff", units="in", width=12.2, height=8.5, res=300)
q <- ggplot(Y_combined, aes(x=Strain, y=`V Counts`, fill=Strain)) +
      geom_boxplot(outlier.shape = NA, width = 0.7, coef=1, varwidth=F, show.legend = F, s
ize=0.9, position = position_dodge(0.9)) +
      geom_jitter(color = "darkgray", alpha =0.6, size=3.3, show.legend = F, position = po
sition_jitterdodge(dodge.width=1.2))+
   facet_wrap(~Antigen, ncol=5, scales="free_x") +
      theme_minimal() +
      chart_design +
      ylab("Log Normalized Intensity")  +
   scale_x_discrete(labels= c("BALB/c", "NOD", "NOR")) +
      labs(title=factor(Y_combined$Antigen), hjust=0.5) +
      scale_fill_jco()
print(q)
dev.off()
```

```
## quartz_off_screen
##                 2
```

```
(q)
```

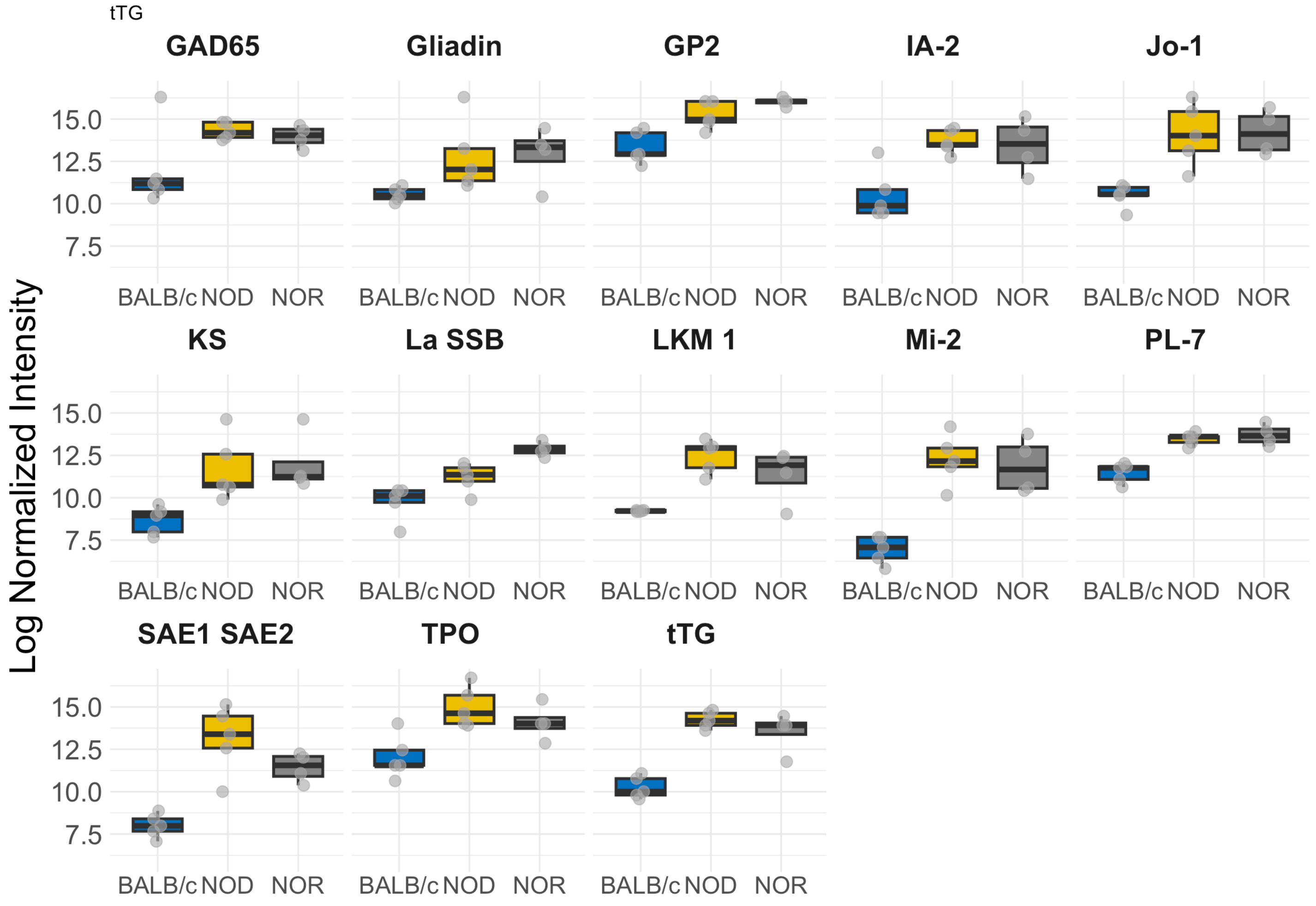


#### Code for Comparing Tear IgG and IgA

```
Tear_IgAhits <- read.csv("IgA_Tears_Voom_Hits.csv", header=T)
Tear_IgGhits <- read.csv("IgG_Tears_Voom_Hits.csv", header=T)

Tear_Ig <- full_join(Tear_IgAhits[,c(1,5,6,7,8)], Tear_IgGhits[,c(1,5,6,7,8)], by="X")

write.csv(Tear_Ig,file="IgA_IgG_Overlap_Tears_Hits.csv", sep=',')
```

##### Total tear IgG total IgA intensity values.

```
Tear_IgG_NSI_raw <- read.csv("~/Documents/3_Parkinsons_disease/Autoantibody_Data/Tear_Auto
_Validation_2022/IgG_MCF_SSK_546_Tear_NSI.csv", header=TRUE, row.names = 1, nrows=88)
Tear_IgG_NSI_raw <- Tear_IgG_NSI_raw[,1:15]
Tear_IgG_SNR_raw <- read.csv("~/Documents/3_Parkinsons_disease/Autoantibody_Data/Tear_Auto
_Validation_2022/IgG_MCF_SSK_546_Tear_SNR.csv", header=TRUE, row.names = 1, , nrows=88)

Tear_IgA_NSI_raw <- read.csv("~/Documents/3_Parkinsons_disease/Autoantibody_Data/Tear_Auto
_Validation_2022/IgA_data/IgA_MCF_SSK_546_Tear_NSI.csv", header=TRUE, row.names = 1)
Tear_IgA_NSI_raw <- Tear_IgA_NSI_raw[1:88,]

x = colSums(Tear_IgG_NSI_raw[1:80,1:14])
y = colSums(Tear_IgA_NSI_raw[1:80,1:14])

data <- as.data.frame(cbind(x,y))
colnames(data) <- c("IgG","IgA")
Strain <- c(rep("NOD", each=5),rep("NOR", each=4), rep("BALBc", each=5))
data$Strain <- factor(Strain)
data$Sample <- rownames(data)
data <- gather(data, "Ig", "Intensity", 1:2)

chart_design <- theme(
   #plot.title = element_text(color = "Black", size = 16, face = "bold", margin = margin(b
=15), hjust=0.4),,
   axis.text.x = element_text(size=16, margin = margin(b=5)),
   axis.text.y = element_text(size=18),
   axis.title.x = element_blank(),
   legend.text = element_text(size=10),
   legend.title = element_blank(),
   legend.position = "top",
   axis.title.y = element_text(size=20, margin = margin(r = 5)),
   strip.text.x = element_text(size =19, margin = margin(b=15), face='bold', hjust=0.4),
   strip.background = element_blank(),
   strip.placement = "outside")

p <- ggplot(data, aes(x=Strain, y=(Intensity), fill=Ig)) +
   geom_boxplot(outlier.shape = NA, width = 0.8, coef=1, varwidth=F, show.legend = T, size
=0.9, position = position_dodge(0.9)) +
   geom_jitter(aes(colour = Ig), alpha =0.6, size=2.5, show.legend = T, position = positio
n_jitterdodge(dodge.width=0.9)) +
   scale_fill_jco() +
   theme_minimal() +
   ylab("Total Intensity of Autoantibodies") +
   chart_design #+
   #ylim(1,(290000))

tiff("ColSum_IgA_vs_IgG.tiff", units="in", width=5.5, height=4.5, res=300)
setwd("~/Documents/3_Parkinsons_disease/Autoantibody_Data/Tear_Auto_Validation_2022/")
print(p)
dev.off()
```

```
## quartz_off_screen
##                 2
```

```
print(p)
```

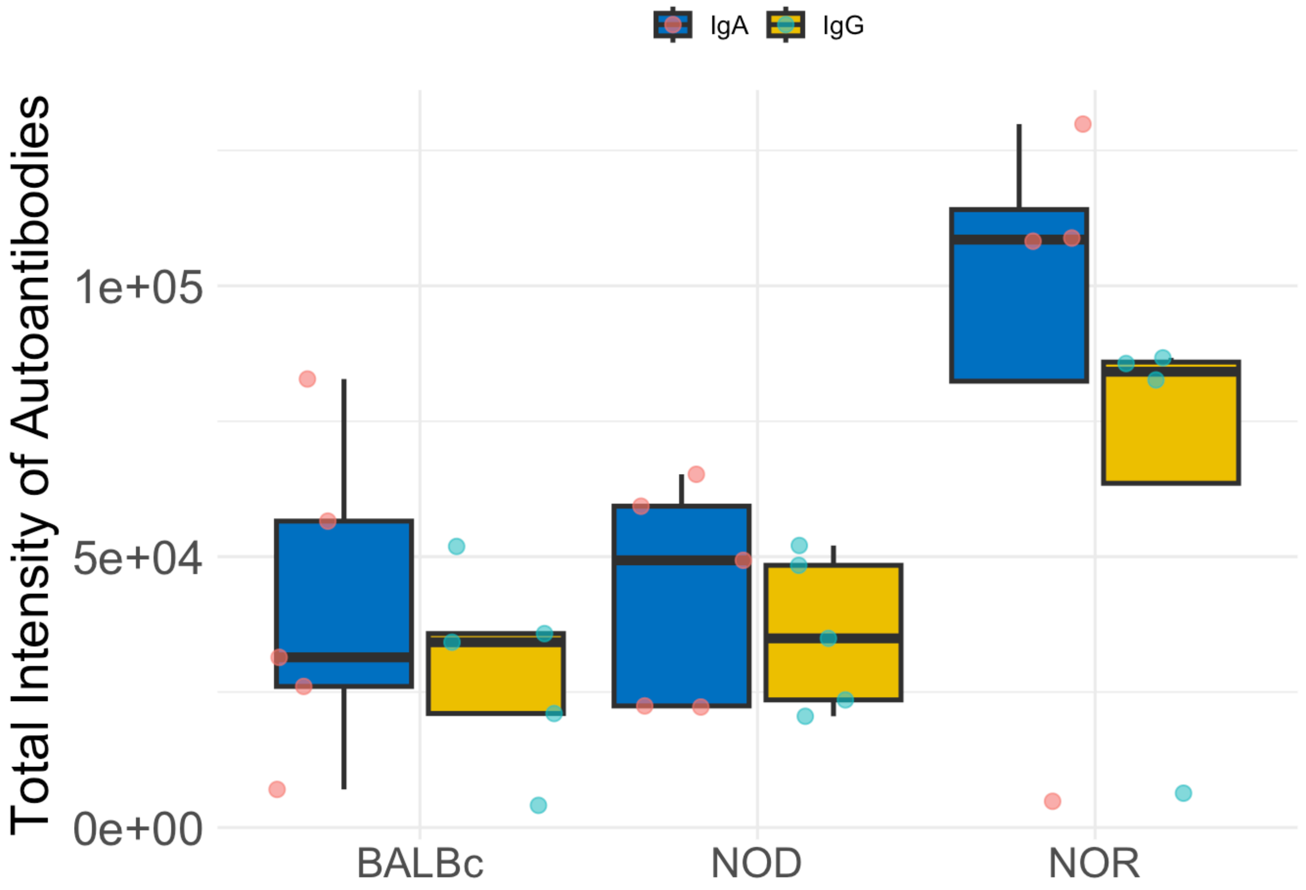


While sum of intensity of 80 autoantigens for IgA is higher in NOR mice tears than IgG, this increase is not statistically significant.

No significant difference was observed between the mean signal IgG intensity as compared to IgA signal intensity for the 80 autoantigens tested in either male NOD or BALB/c mice.

```
mod <- lm((Intensity) ~ Strain*Ig, data)
aov(mod)
```

```
## Call:
##    aov(formula = mod)
##
## Terms:
##                      Strain         Ig   Strain:Ig   Residuals
## Sum of Squares   8888312269 1239835830   258090631 21357553132
## Deg. of Freedom           2          1           2          22
##
## Residual standard error: 31157.63
## Estimated effects may be unbalanced
```

```
summary(aov(mod))
```

```
##             Df    Sum Sq   Mean Sq F value Pr(>F)
## Strain       2 8.888e+09 4.444e+09   4.578 0.0218 *
## Ig           1 1.240e+09 1.240e+09   1.277 0.2706
## Strain:Ig    2 2.581e+08 1.290e+08   0.133 0.8762
## Residuals   22 2.136e+10 9.708e+08
## ---
## Signif. codes:  0 '***' 0.001 '**' 0.01 '*' 0.05 '.' 0.1 ' ' 1
```

```
TukeyHSD(aov(mod))
```

```
##   Tukey multiple comparisons of means
##     95% family-wise confidence level
##
## Fit: aov(formula = mod)
##
## $Strain
##                diff        lwr      upr     p adj
## NOD-BALBc  4713.39 -30289.988 39716.77 0.9390493
## NOR-BALBc 41548.73   4422.041 78675.42 0.0264323
## NOR-NOD   36835.34   -291.349 73962.03 0.0520810
##
## $Ig
##              diff       lwr     upr     p adj
## IgG-IgA -13308.62 -37731.54 11114.3 0.2706042
##
## $`Strain:Ig`
##                          diff        lwr        upr     p adj
## NOD:IgA-BALBc:IgA     2936.78  -58449.53  64323.095 0.9999881
## NOR:IgA-BALBc:IgA    47172.26  -17937.75 112282.284 0.2530846
## BALBc:IgG-BALBc:IgA -11364.18  -72750.49  50022.135 0.9915938
## NOD:IgG-BALBc:IgA    -4874.18  -66260.49  56512.135 0.9998539
## NOR:IgG-BALBc:IgA    24561.01  -40549.00  89671.034 0.8437306
## NOR:IgA-NOD:IgA      44235.48  -20874.53 109345.504 0.3153347
## BALBc:IgG-NOD:IgA   -14300.96  -75687.27  47085.355 0.9765795
## NOD:IgG-NOD:IgA      -7810.96  -69197.27  53575.355 0.9985525
## NOR:IgG-NOD:IgA      21624.23  -43485.78  86734.254 0.9010848
## BALBc:IgG-NOR:IgA   -58536.44 -123646.46   6573.574 0.0948797
## NOD:IgG-NOR:IgA     -52046.44 -117156.46  13063.574 0.1700316
## NOR:IgG-NOR:IgA     -22611.25  -91243.24  46020.737 0.9040227
## NOD:IgG-BALBc:IgG     6490.00  -54896.31  67876.315 0.9994078
## NOR:IgG-BALBc:IgG    35925.19  -29184.82 101035.214 0.5344782
## NOR:IgG-NOD:IgG      29435.19  -35674.82  94545.214 0.7218548
```

```
plot(mod, which = 2)
```

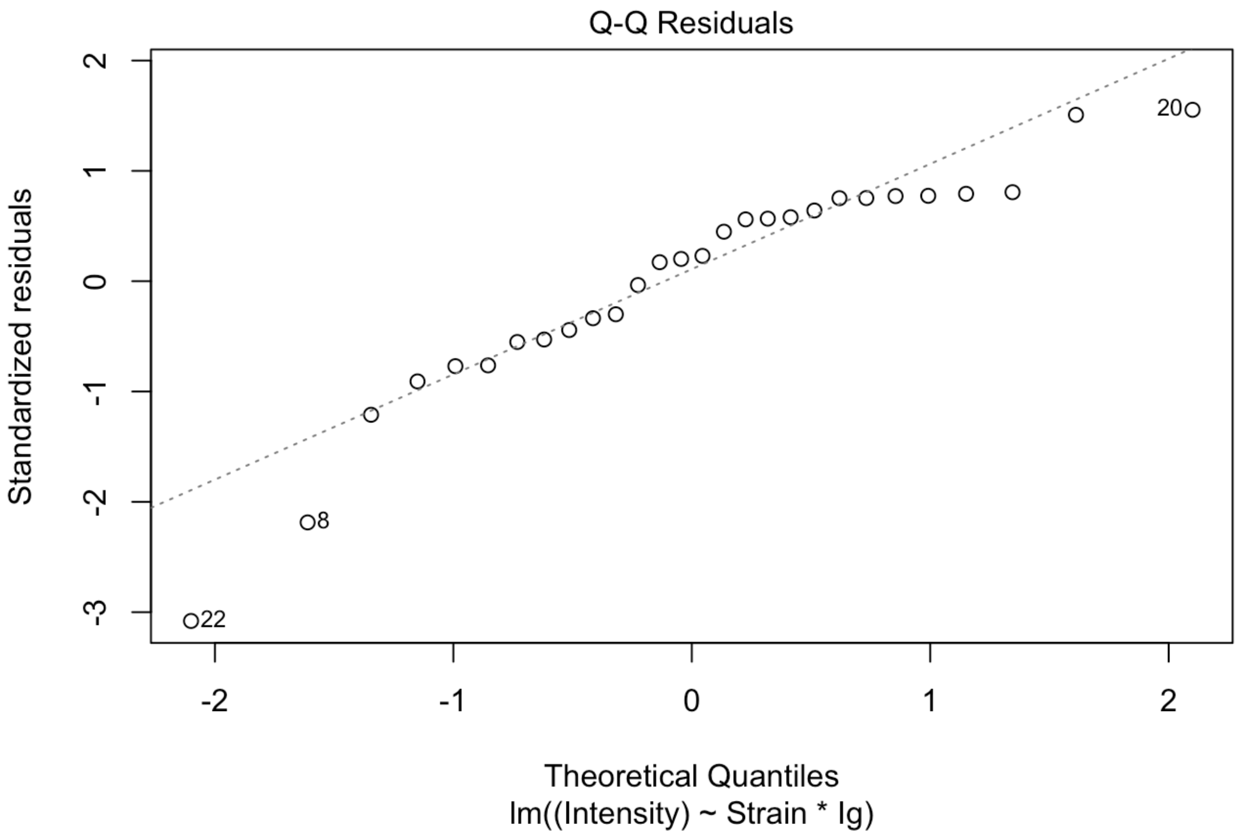


```
hist(mod$residuals)
```

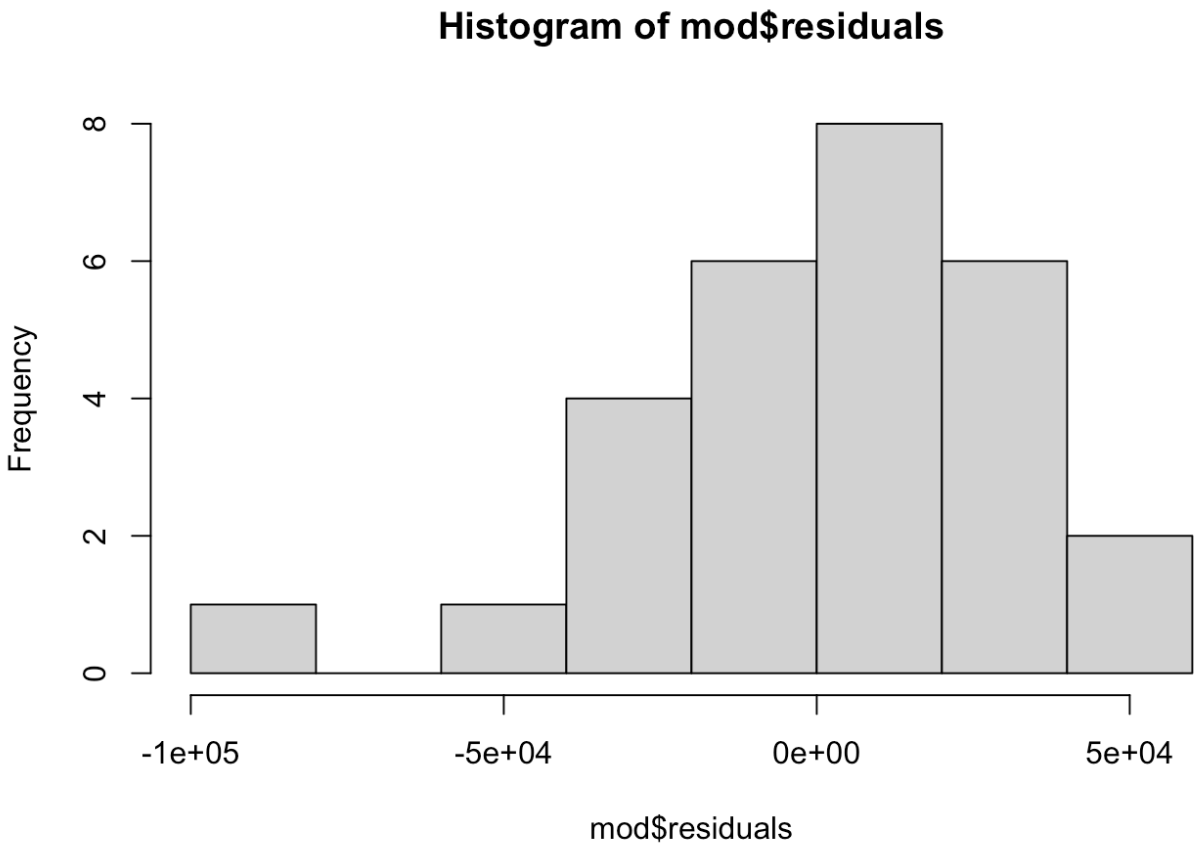


```
shapiro.test(mod$residuals) #data are non-normal
```

```
##
##  Shapiro-Wilk normality test
##
## data:  mod$residuals
## W = 0.90997, p-value = 0.01974
```

```
plot(mod, which = 3)
```

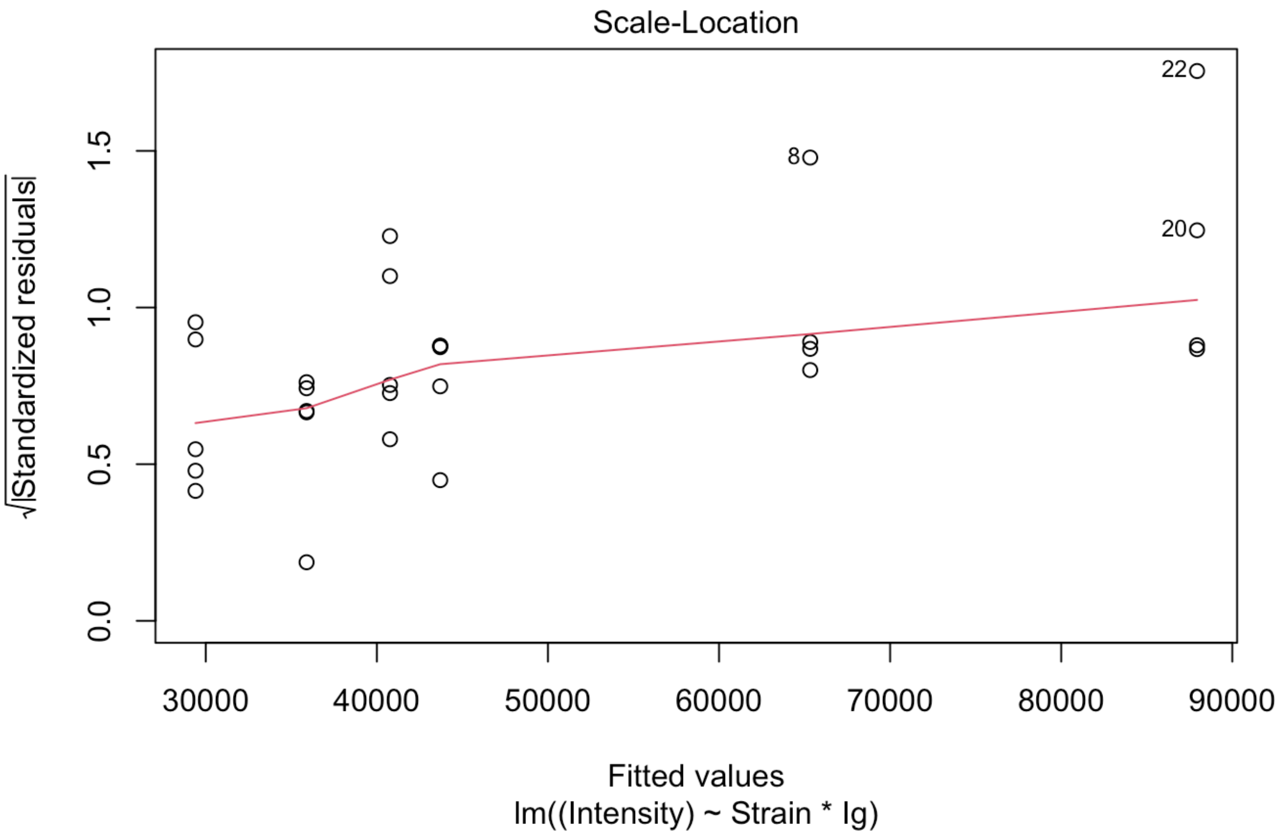


```
#leveneTest(aov(Intensity ~ Strain + Ig, data=data))
```

#### Chunk for plotting IgG vs IgA secondary control for each of the samples

```
#x1 = unlist(Tear_IgG_NSI_raw[87,2:16])
#x2 = unlist(Tear_IgG_NSI_raw[88,2:16])
#y1 = unlist(Tear_IgA_NSI_raw[87,])
#y2 = unlist(Tear_IgA_NSI_raw[88,])
Strain <- c(rep("NOD", each=5),rep("NOR", each=4), rep("BALBc", each=6))
x = unlist(lapply(Tear_IgG_NSI_raw[87:88,], mean, 2))
y = unlist(lapply(Tear_IgA_NSI_raw[87:88,], mean, 2))
data <- as.data.frame(cbind(x,y))
colnames(data) <- c("IgG","IgA")
data$Strain <- factor(Strain)
data$Sample <- rownames(data)
data <- gather(data, "Ig", "Intensity", 1:2)

tiff("Tear_IgA_vs_IgG.tiff", units="in", width=5.5, height=4.5, res=300)
setwd("~/Documents/3_Parkinsons_disease/Autoantibody_Data/Tear_Auto_Validation_2022/")
q <- ggplot(data, aes(x=Strain, y=(Intensity), fill=Ig)) +
   geom_boxplot(outlier.shape = NA, width = 0.8, coef=1, varwidth=F, show.legend = T, size
=0.9, position = position_dodge(0.9)) +
   geom_jitter(aes(colour = Ig), alpha =0.6, size=2.5, show.legend = T, position = positio
n_jitterdodge(dodge.width=0.9)) +
   scale_fill_jco() +
   theme_minimal() +
   ylab("Average Intensity of Ig Controls") +
   chart_design #+
   #ylim(0,100000)
print(q)
dev.off()
```

```
## quartz_off_screen
##                 2
```

```
print(q)
```

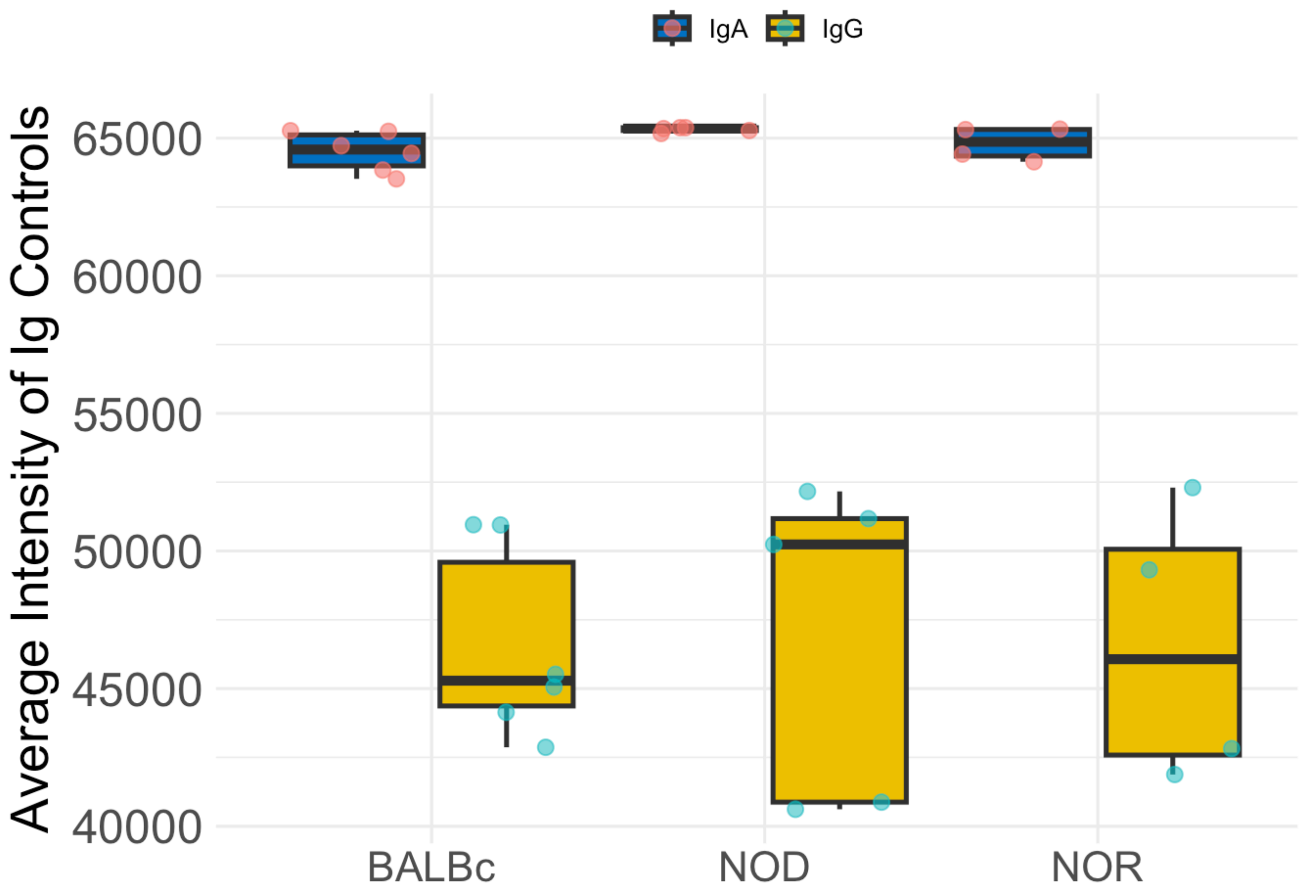


```
#+ ylim(10,12)
```

Signal intensity observed for mouse IgA secondary control was significantly higher than the intensity observed for mouse IgG secondary control in the tears of male NOD, NOR and BALB/c mice. Levels of signal intensity for either IgG or IgA were consistent between the three mouse groups.

```
#Statistics
mod <- lm((Intensity) ~ Strain*Ig, data)
aov(mod)
```

```
## Call:
##    aov(formula = mod)
##
## Terms:
##                    Strain         Ig  Strain:Ig  Residuals
## Sum of Squares    2188792 2464619432     208842  274448157
## Deg. of Freedom         2          1          2         24
##
## Residual standard error: 3381.618
## Estimated effects may be unbalanced
```

```
summary(aov(mod))
```

```
##             Df    Sum Sq   Mean Sq F value   Pr(>F)    
## Strain       2 2.189e+06 1.094e+06   0.096    0.909    
## Ig           1 2.465e+09 2.465e+09 215.527 1.73e-13 ***
## Strain:Ig    2 2.088e+05 1.044e+05   0.009    0.991    
## Residuals   24 2.744e+08 1.144e+07                     
## ---
## Signif. codes:  0 '***' 0.001 '**' 0.01 '*' 0.05 '.' 0.1 ' ' 1
```

```
TukeyHSD(aov(mod))
```

```
##   Tukey multiple comparisons of means
##     95% family-wise confidence level
## 
## Fit: aov(formula = mod)
## 
## $Strain
##                  diff       lwr      upr     p adj
## NOD-BALBc  617.6833 -2998.194 4233.561 0.9049489
## NOR-BALBc  145.2708 -3709.268 3999.809 0.9951292
## NOR-NOD   -472.4125 -4478.166 3533.341 0.9534036
## 
## $Ig
##              diff       lwr       upr p adj
## IgG-IgA -18127.77 -20676.25 -15579.28     0
## 
## $`Strain:Ig`
##                             diff        lwr        upr     p adj
## NOD:IgA-BALBc:IgA      802.2917  -5528.967   7133.550 0.9986460
## NOR:IgA-BALBc:IgA      291.7292  -6457.415   7040.874 0.9999932
## BALBc:IgG-BALBc:IgA -17926.5833 -23963.202 -11889.965 0.0000000
## NOD:IgG-BALBc:IgA   -17493.5083 -23824.767 -11162.250 0.0000001
## NOR:IgG-BALBc:IgA   -17927.7708 -24676.915 -11178.626 0.0000003
## NOR:IgA-NOD:IgA       -510.5625  -7524.479   6503.354 0.9999093
## BALBc:IgG-NOD:IgA   -18728.8750 -25060.134 -12397.616 0.0000000
## NOD:IgG-NOD:IgA     -18295.8000 -24908.584 -11683.016 0.0000001
## NOR:IgG-NOD:IgA     -18730.0625 -25743.979 -11716.146 0.0000003
## BALBc:IgG-NOR:IgA   -18218.3125 -24967.457 -11469.168 0.0000002
## NOD:IgG-NOR:IgA     -17785.2375 -24799.154 -10771.321 0.0000006
## NOR:IgG-NOR:IgA     -18219.5000 -25612.817 -10826.183 0.0000010
## NOD:IgG-BALBc:IgG      433.0750  -5898.184   6764.334 0.9999333
## NOR:IgG-BALBc:IgG       -1.1875  -6750.332   6747.957 1.0000000
## NOR:IgG-NOD:IgG       -434.2625  -7448.179   6579.654 0.9999593
```

```
plot(mod, which = 2)
```

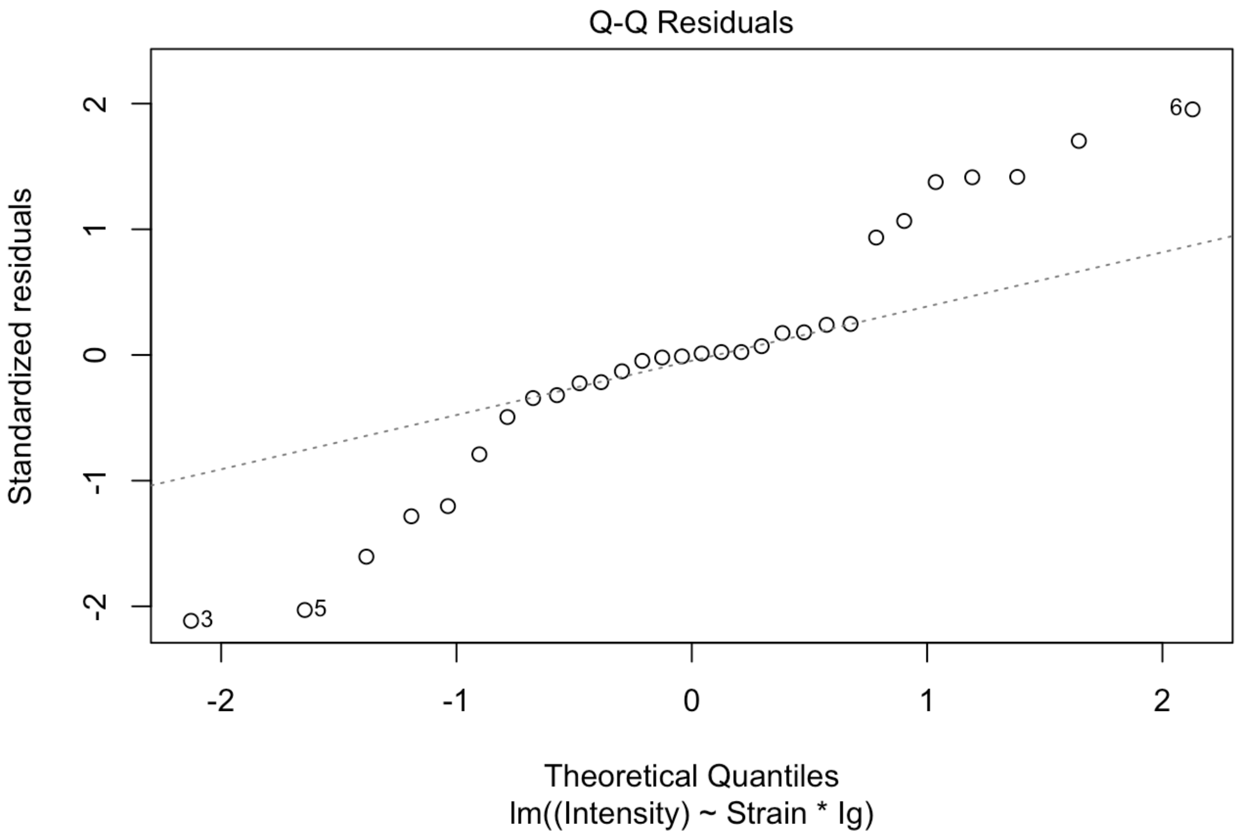


```
hist(mod$residuals)
```

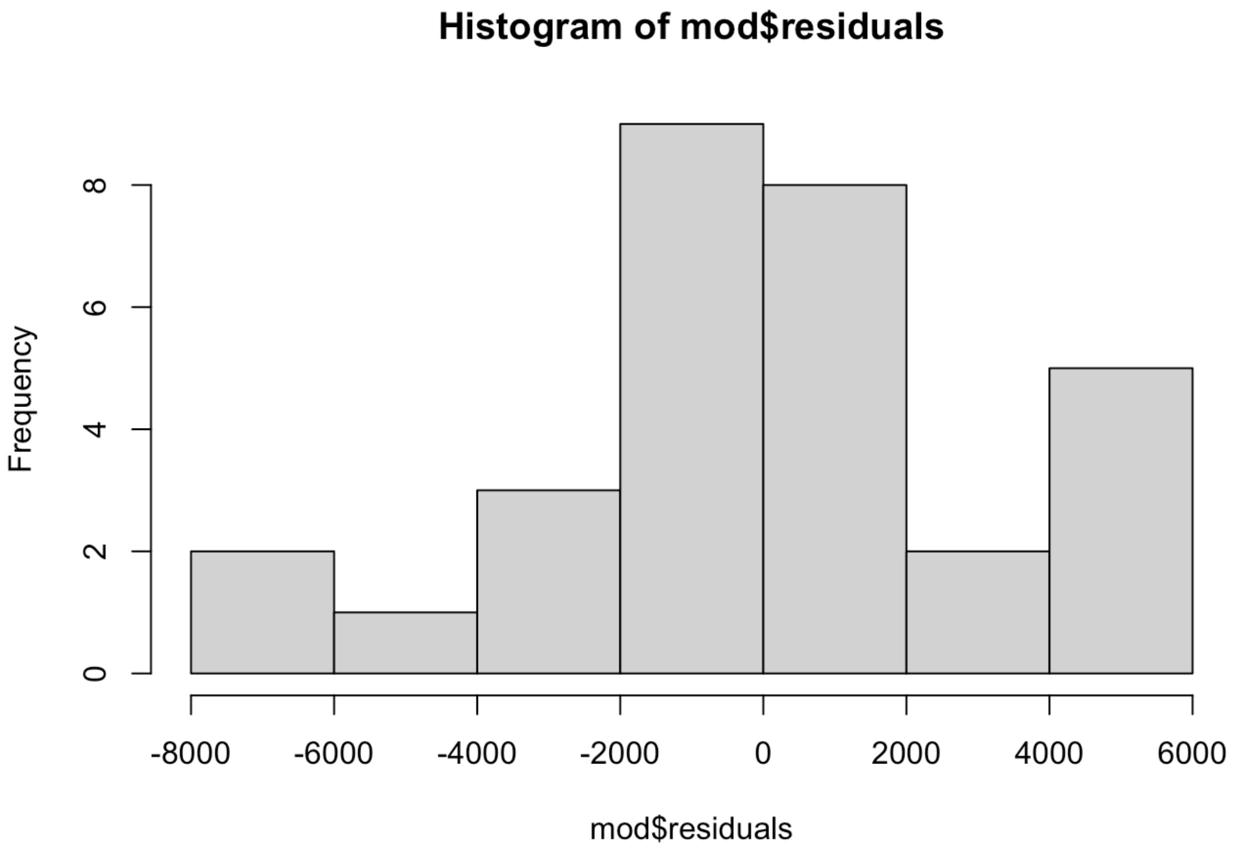


```
shapiro.test(mod$residuals) #data are normally distributed
```

```
##
##  Shapiro-Wilk normality test
##
## data:  mod$residuals
## W = 0.94833, p-value = 0.1525
```

```
plot(mod, which = 3)
```

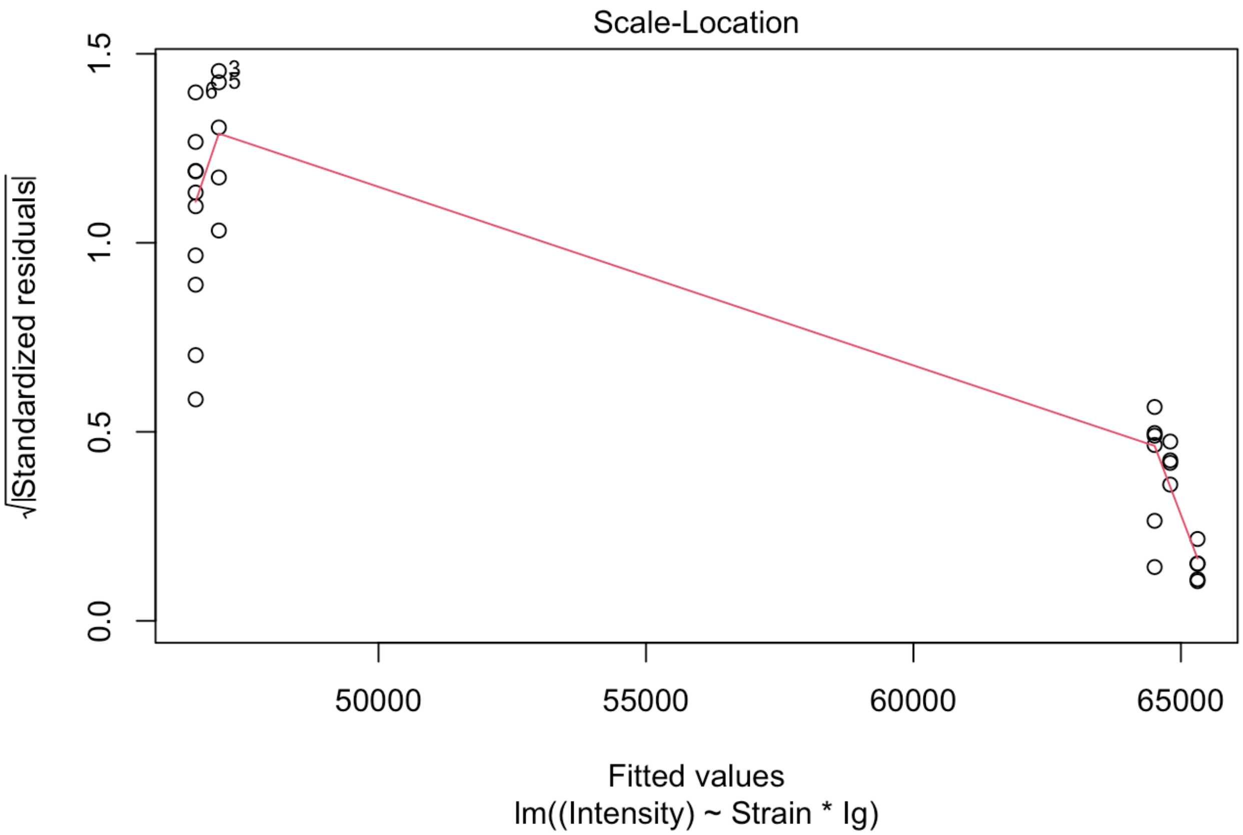


### Tear IgG vs IgA Correlation Analysis

Scatterplots showing raw signal intensity for IgG vs IgA of each autoantigen in the tears of NOD, NOR and BALB/c mice.

```
#which((rowSums(Tear_IgG_SNR_raw[1:80,1:15])/15) < 2.8)
#which((rowSums(IgA_SNR[1:80,])/15) < 2.8)

Tear_IgA_NSI <- Tear_IgA_NSI_raw[1:80,]
Tear_IgA_NSI <- Tear_IgA_NSI[-c(which((rowSums(Tear_IgG_SNR_raw[1:80,1:15])/15) < 2.8 & (r
owSums(IgA_SNR[1:80,])/15) < 2.8)),]
Tear_IgA_NSI$Antigen <- rownames(Tear_IgA_NSI)
Tear_IgA_NSI <- gather(Tear_IgA_NSI, "Sample", "Intensity", 1:(ncol(Tear_IgA_NSI)-1))
Tear_IgA_NSI$Strain <- str_sub(Tear_IgA_NSI$Sample, 1, (str_locate(Tear_IgA_NSI$Sample,
"_")[,1] -1))
Tear_IgA_NSI$Ig <- "IgA"

Tear_IgG_NSI <- Tear_IgG_NSI_raw[1:80,]
Tear_IgG_NSI <- Tear_IgG_NSI[-c(which((rowSums(Tear_IgG_SNR_raw[1:80,])/15) < 2.8 & (rowSu
ms(IgA_SNR[1:80,])/15) < 2.8)),]
Tear_IgG_NSI$Antigen <- rownames(Tear_IgG_NSI)
Tear_IgG_NSI <- gather(Tear_IgG_NSI, "Sample", "Intensity", 1:(ncol(Tear_IgG_NSI)-1))
Tear_IgG_NSI$Strain <- str_sub(Tear_IgG_NSI$Sample, 1, (str_locate(Tear_IgG_NSI$Sample,
"_")[,1] -1))
Tear_IgG_NSI$Ig <- "IgG"

data_mat <- dplyr::bind_rows(Tear_IgG_NSI, Tear_IgA_NSI)

###Plot for NODs & NORs
data_mat2 <- dplyr::full_join(Tear_IgA_NSI, Tear_IgG_NSI, by = c("Antigen", "Sample"),suff
ix = c(".IgA", ".IgG"),)

### Plot for NORs
ggplot(data = data_mat2[which(data_mat2$Strain.IgA=="NOR"),]) +
   geom_point(aes(x=(Intensity.IgA), y=(Intensity.IgG), fill=factor(Antigen), color=factor
(Antigen)), show.legend = F) +
   facet_wrap(~factor(Antigen))
```

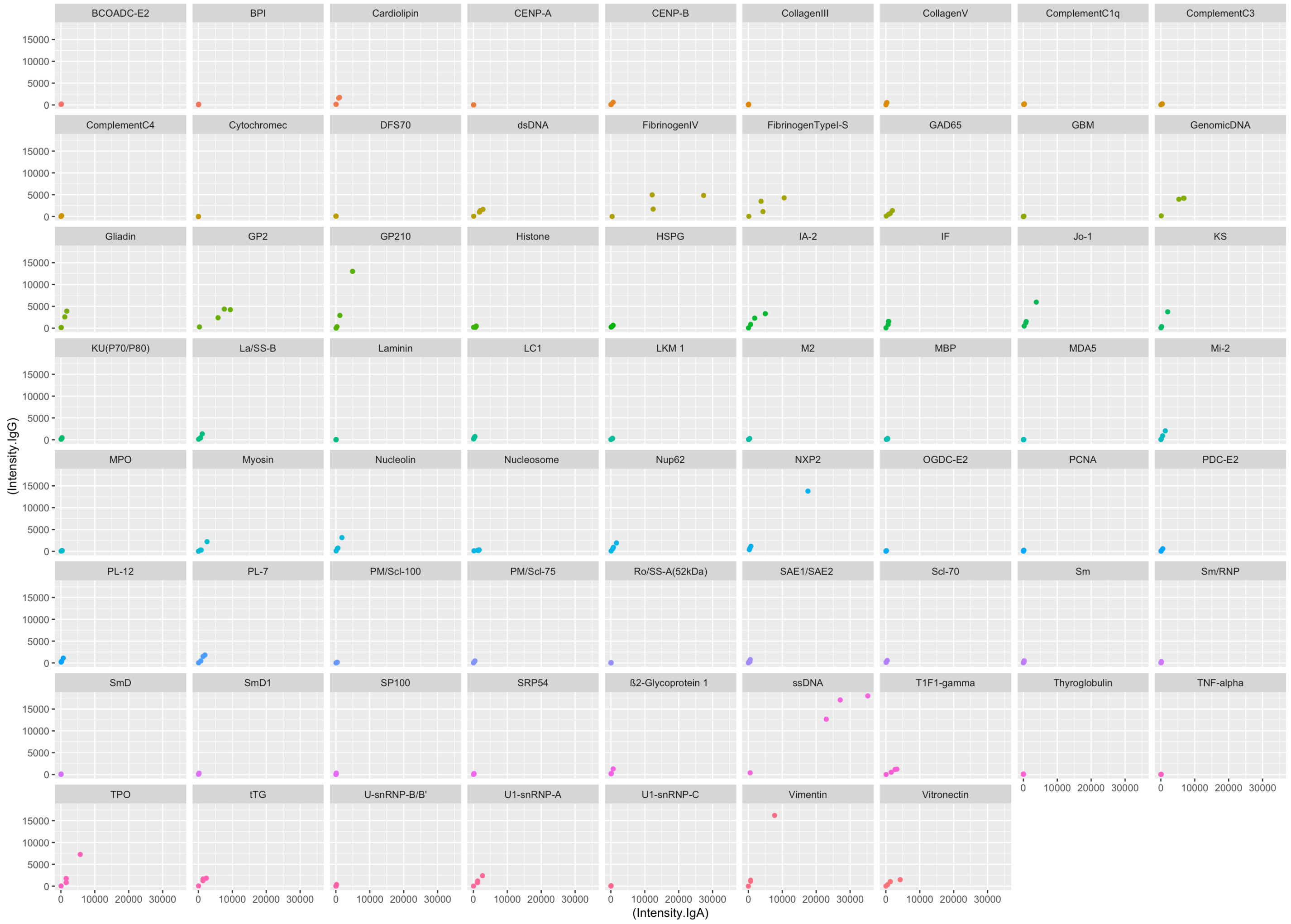


There is a need to normalize the x and y axes. In the next plot, we run correlation on log10 normalized NSI for IgG and IgA.

### Correlation Analysis Figure

Sctterplots of IgG vs IgA for NOD, NOR and BALB/c. Each point represents one mouse. Spearman correlation coefficient rho and p-values are shown. p < 0.01 is considered significant.

```
### Plot for NOD, NOR & BALB/c
data_mat2$model <- ifelse(data_mat2$Strain.IgA=="BALBc", "Healthy", "SjD")

p <- ggplot(data = data_mat2, aes(x=log10(Intensity.IgA+1), y=log10(Intensity.IgG+1))) +
   geom_point(aes( color=factor(model), fill=factor(Antigen)), show.legend = F, alpha = 0.
8) +
   geom_smooth(aes(color = factor(model)),method=lm, se=FALSE, fullrange=FALSE, linewidth=
0.25) +
   scale_color_jco(guide="legend") +
   guides(fill="none") +
   theme_minimal() + ylim(-1, 4.6) +
   xlab("Log 10 IgA NSI") +
   ylab("Log 10 IgG NSI") +
   stat_cor(aes(color=factor(model)), method = "spearman", r.digits=2, p.accuracy=0.0005,
label.y = c(-0.9,4.4), cor.coef.name="rho") +
   facet_wrap(~factor(Antigen),nrow=7) +
   chart_design

print(p)
```

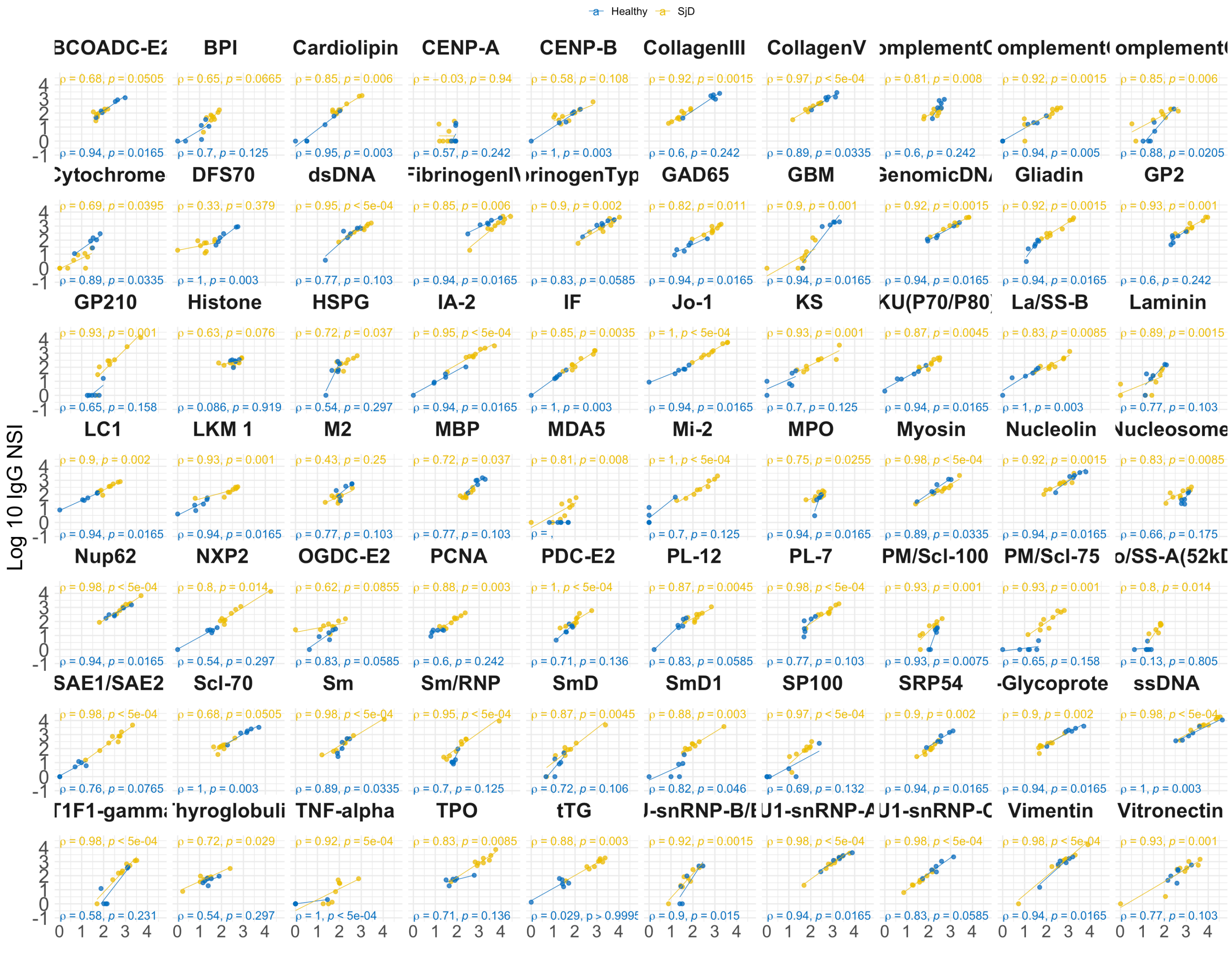

```
setwd("~/Documents/3_Parkinsons_disease/Autoantibody_Data/Tear_Auto_Validation_2022")
tiff("All_IgGvsIgA_Tears.jpeg", units="in", width=16, height=10, res=300)
print(p)
dev.off()
```

```
## quartz_off_screen
##                 2
```

#### Scatterplot of shared epitopes between upregulated IgG and IgA autoantibodies

Scatterplot of Log10 IgG vs Log10 IgA NSI of shares epitopes of autoantibodies that are significantly upregulated in male NOD and male NOR mice tears when compared to BALB/c controls. NOD and NOR are shown in yellow (referred to an SjD) and BALB/c are shown in blue (referred to as Healthy).

```
data_mat3 <- data_mat2[which(data_mat2$Antigen %in% c("tTG", "TPO", "Mi-2", "Jo-1", "IA-
2", "SAE1/SAE2 ")),]
p <- ggplot(data = data_mat3, aes(x=log10(Intensity.IgA+1), y=log10(Intensity.IgG+1))) +
   geom_point(aes( color=factor(model), fill=factor(Antigen)), show.legend = T, alpha = 0.
8) +
   geom_smooth(aes(color = factor(model)),method=lm, se=FALSE, fullrange=FALSE, linewidth=
0.25) +
   scale_color_jco(guide="legend") +
   guides(fill="none") +
   theme_minimal() + ylim(-1, 4.6) +
   xlab("Log 10 IgA NSI") +
   ylab("Log 10 IgG NSI") +
   stat_cor(aes(color=factor(model)), method = "spearman", r.digits=2, p.accuracy=0.0005,
label.y = c(-0.9,4.4), cor.coef.name="rho") +
   facet_wrap(~factor(Antigen),nrow=2) +
   chart_design

print(p)
```

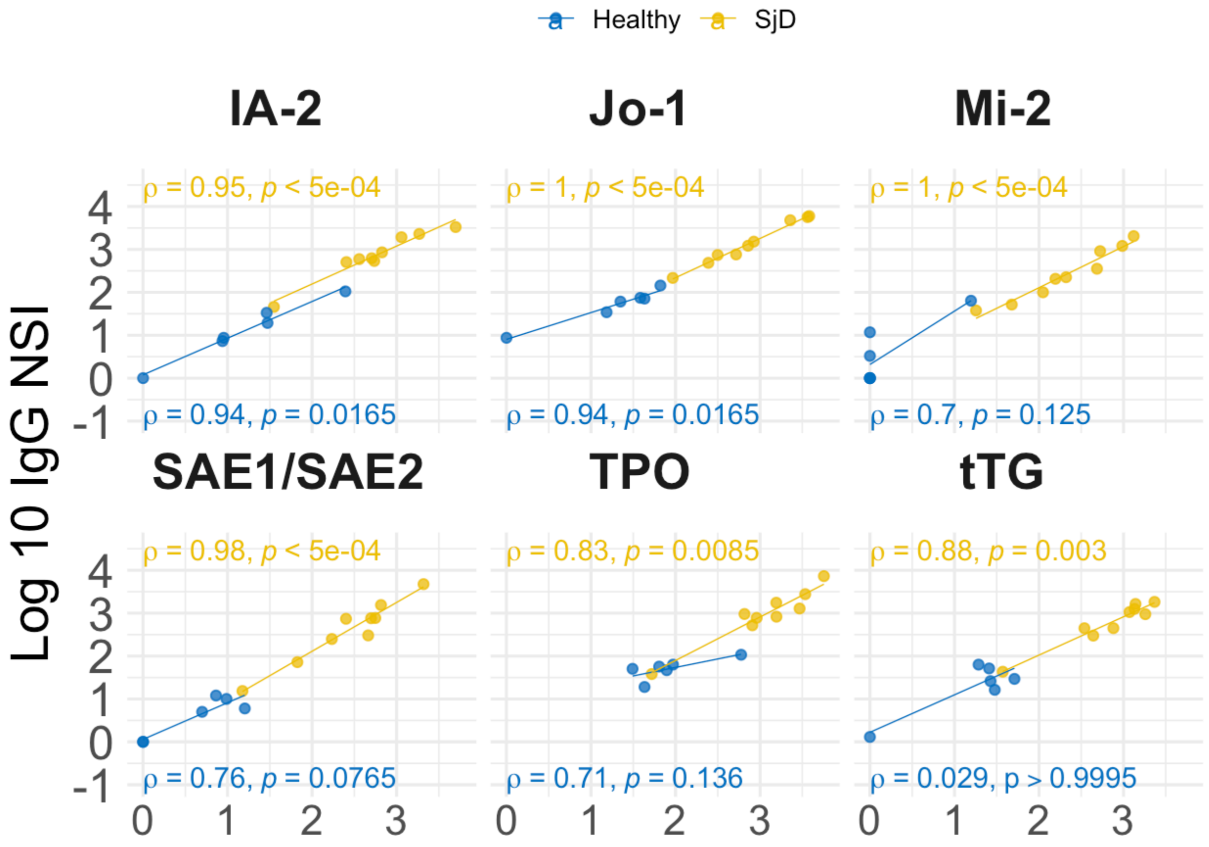


```
setwd("~/Documents/3_Parkinsons_disease/Autoantibody_Data/Tear_Auto_Validation_2022")
tiff("Supp_Figure_6.jpeg", units="in", width=6.5, height=4.8, res=300)
print(p)
dev.off()
```

```
## quartz_off_screen
##                 2
```
